# EquiScore: A generic protein-ligand interaction scoring method integrating physical prior knowledge with data augmentation modeling

**DOI:** 10.1101/2023.06.18.545464

**Authors:** Duanhua Cao, Geng Chen, Jiaxin Jiang, Jie Yu, Runze Zhang, Mingan Chen, Wei Zhang, Lifan Chen, Feisheng Zhong, Yingying Zhang, Chenghao Lu, Xutong Li, Xiaomin Luo, Sulin Zhang, Mingyue Zheng

**Author notes:** **Corresponding Authors:** (Mingyue Zheng). **Author Contributions:** D.H.C., G.C. contributed equally to this study. M.Y.Z. designed the research study. D.H.C developed the method and implemented the code. G.C., collected and processed training data. D.H.C, G.C, J.X.J., D.H.C, and J.Y. benchmarked the methods. All authors contributed to the analysis of the results D.H.C., G. C. and M.Y.Z. wrote the paper. All authors read and approved the manuscript. **Notes:** The authors declare no competing financial interest.

## Abstract

Developing robust methods for evaluating protein-ligand interactions has been a long-standing problem. Here, we propose a novel approach called EquiScore, which utilizes an equivariant heterogeneous graph neural network to integrate physical prior knowledge and characterize protein-ligand interactions in equivariant geometric space. To improve generalization performance, we constructed a dataset called PDBscreen and designed multiple data augmentation strategies suitable for training scoring methods. We also analyzed potential risks of data leakage in commonly used data-driven modeling processes and proposed a more stringent redundancy removal scheme to alleviate this problem. On two large external test sets, EquiScore outperformed 21 methods across a range of screening performance metrics, and this performance was insensitive to binding pose generation methods. EquiScore also showed good performance on the activity ranking task of a series of structural analogs, indicating its potential to guide lead compound optimization. Finally, we investigated different levels of interpretability of EquiScore, which may provide more insights into structure-based drug design.

## INTRODUCTION

After the Human Genome Project, the challenge of translating new knowledge from genomics into new medicines has arisen. In recent years, there have been breakthroughs in protein folding algorithms, resulting in dramatic progress in the field of structural biology^1, 2^. An ambitious project has been proposed to find specific ligands or probes for the entire human proteome^3^. Once a high-quality protein structure is available, we can use structure-based virtual screening (SBVS) to select only the best-fitting molecules for synthesis and testing. For example, molecular docking approaches can be used to explore large chemical space. These approaches are gaining renewed attention due to the growing availability of many bespoke or make-on-demand virtual libraries^4, 5^. While significant progress has been made in this field, developing a scoring method with higher accuracy in practical application scenarios remains an open challenge^6–8^.

The scoring method based on machine learning has made significant progress with the explosive growth of experimental protein-ligand interaction data. Various machine learning algorithms and neural network architectures, such as three-dimensional convolutional neural networks (3D-CNNs)^9, 10^, and graph convolutional neural networks (GNNs)^11–16^ have shown improvements in screening and scoring power on benchmarks^9–15^. However, the performance of these data-driven models is often system-dependent and difficult to generalize to protein or ligand chemical types that are not included in the model training process. A comparative analysis revealed that machine learning scoring methods do not outperform traditional scoring methods on unseen targets in their training set^18^. This highlights the need for more robust and reliable methods to better address such out-of-distribution (OOD) challenges.

Two factors primarily limit the generalizability of scoring methods: the data used to train the model and the algorithms that learn from the data. PDBbind^19^ and DUD-E^20^ represent the two most commonly used types of datasets. PDBbind contains protein-ligand binding complex structures and associated binding affinity data that can be used to train regression models between the structure and activity. In contrast, DUD-E contains both “real” and “decoy” protein-ligand binding complex structures, which are generally used to train classification models that can distinguish positive and negative samples. Although the first type of dataset is more favorable because the regression method can quantitatively predict binding affinity and have more applicable scenarios, the amount of such association data is limited, and they do not contain negative samples. This can easily lead to a high false positive rate in virtual screening (VS) settings for methods derived from these datasets. The second type of dataset contains more negative samples, and the resulting classification methods may have an advantage in discriminating negative samples and reducing the false positive rate. However, many active compounds in such datasets have similar chemical structures or the same skeleton, resulting in a significant properties distribution bias between positive and negative samples. Many studies have found that machine learning-based scoring methods tend to memorize the inherent biases of the training data rather than learning features of protein-ligand interactions, resulting in limited generalization ability^21–23^.

In summary, problems with training data primarily relate to two aspects. First, positive sample volumes and diversity are often insufficient, resulting in limited information that the model can utilize. Second, many public datasets suffer from internal data distribution biases that may prevent the model from learning the protein-ligand interactions we seek to understand.

Regarding the factor of algorithms, various neural network architectures suitable for solving different types of data problems have been leveraged in developing scoring methods. However, directly applying these architectures to address protein-ligand interaction prediction still has many deficiencies. For instance, 3D-CNNs^9, 10^ require extensive data augmentation to account for equivariance in 3D rotation and translation of atoms. GNNs^11–16^ may ignore some important information in the complex, such as building edges with a specific distance threshold, which loses the prior knowledge of the chemical structure and cannot accurately characterize distance-dependent interatomic physical interactions in the protein-ligand complex. For example, hydrogen bonds and van der Waals interactions are more sensitive to interatomic distance than electrostatic interactions. The frequently used one-hot encoding that indicates whether an atom is aromatic does not reflect well the non-local contribution of an aromatic ring to intermolecular interactions, such as π-π interactions between aromatic ring systems. Introducing physical prior information into the scoring method is another key issue that can help further improve generalization ability^13, 24, 25^. Recently, equivariant models have shown potential for more accurate and efficient predictions of intermolecular interactions^26–29^. This is because they have more expressive operations on important geometric tensor interactions^27^, such as multiple dipoles or hydrogen bonding interactions^30, 31^. Despite these advances, the introduction of physical inductive bias is still not well considered in these models. Therefore, there is a high demand to investigate novel equivariant neural network architectures that can better learn protein-ligand interactions by integrating physical prior knowledge with data-driven modeling.

This study aims to improve the deep learning-based scoring method in two ways. Firstly, we collect more positive samples and use a molecular deep generative model^32^ to generate more deceptive and diverse decoy molecules, to reduce possible biases in constructing a VS training dataset. Secondly, we introduce an equivariant graph neural network that integrates physical prior knowledge into a heterogeneous graph and adopts a new update mechanism to enable better information interaction. We use the designed data set and the heterogeneous graph network to train the final scoring method, named EquiScore. For evaluation, we (1) compare EquiScore with a comprehensive set of newly reported deep learning scoring methods on two external test sets, DUD-E^20^ and DEKOIS2.0^33, 34^, to evaluate its screening power on unseen protein systems; (2) compare EquiScore with a range of different methods on a lead optimization dataset, LeadOpt^35^, to evaluate its activity ranking ability for structural analogs; and (3) use different docking methods to generate binding poses to further evaluate the robustness of EquiScore as a rescoring method. Finally, we analyzed the interpretability of the model to examine whether it learned the key intermolecular interactions we are interested in. This information could provide meaningful clues for rational drug design.

## RESULTS AND DISCUSSION

### Data preparation

A recent study identified three possible biases in constructing VS training datasets: artificial enrichment, analog bias, and false negative bias^36^. Artificial enrichment arises from distinct differences in physical and chemical properties between positive and negative samples, making it easy for the model to distinguish between them. Analog bias occurs when many positive compounds in a dataset have similar chemical structures or the same skeleton, leading to high enrichment performance. False negative bias arises from using positive samples as negative samples during dataset construction. These biases can limit the trained model’s generalization ability and increase the probability of false positives. Therefore, minimizing the occurrence of these biases when constructing a dataset is a key challenge.

Accordingly, we improved the construction of datasets for the training scoring method in three ways, as shown in the schematic diagram **Fig.1** (Refer to the method section for details). First, we collected complex crystal structures from the PDB database to increase the diversity of positive samples and alleviate the dataset’s analog bias problem. Second, we retained the near-native poses, i.e., with Root-Mean Square Deviation (RMSD) less than 2Å to crystal pose, after re-docking and the pose with the highest docking score as additional positive samples. This procedure aims to introduce noises generated by pose generation methods to increase the model’s generalization ability. Third, for negative sample construction, we first constructed negative samples by cross-docking to ensure that each ligand appears in both positive and negative samples, which we called” label reversal” experiment^37^. This way, the model cannot distinguish positive and negative samples simply by remembering the ligand substructures, and will be forced to learn more difficult higher-level protein-ligand interaction information. To further limit the artificial enrichment bias in the dataset, we generated 500 decoys with similar physical and chemical properties to the ligands of each complex using the generative model DeepCoy^38^. The resulting samples were then docked and clustered by the Shape Screening module^39^ in Schrödinger (Schrödinger, LLC, New York, NY, 2020). We only kept the top 5 decoys whose shape is closest to the crystal ligand pose as negative samples, which can further increase the difficulty of the model for correct recognition, thus alleviating the artificial enrichment bias. The above data augmentation strategies aim to help the model learn representations that can generalize across proteins.

**Fig.1.**
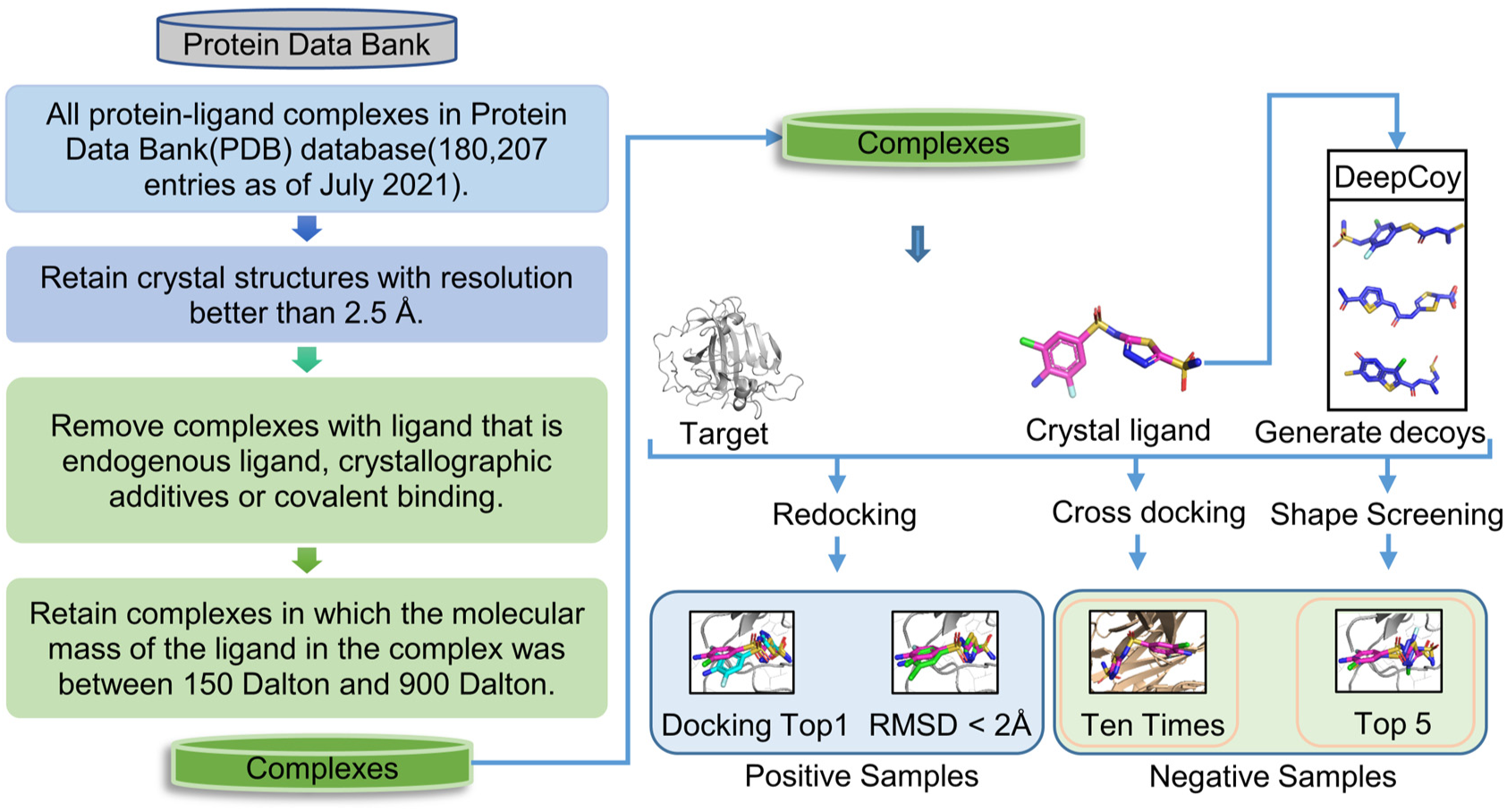
The pipeline of collecting complex data from the PDB database and data augmentation strategies.

Finally, we named the resulting dataset PDBscreen, and its statistics are shown in **Table 1**. In contrast to PDBbind, this dataset includes more crystal complexes without ligand binding affinity data, as well as samples generated through data augmentation strategies.

**Table 1.**
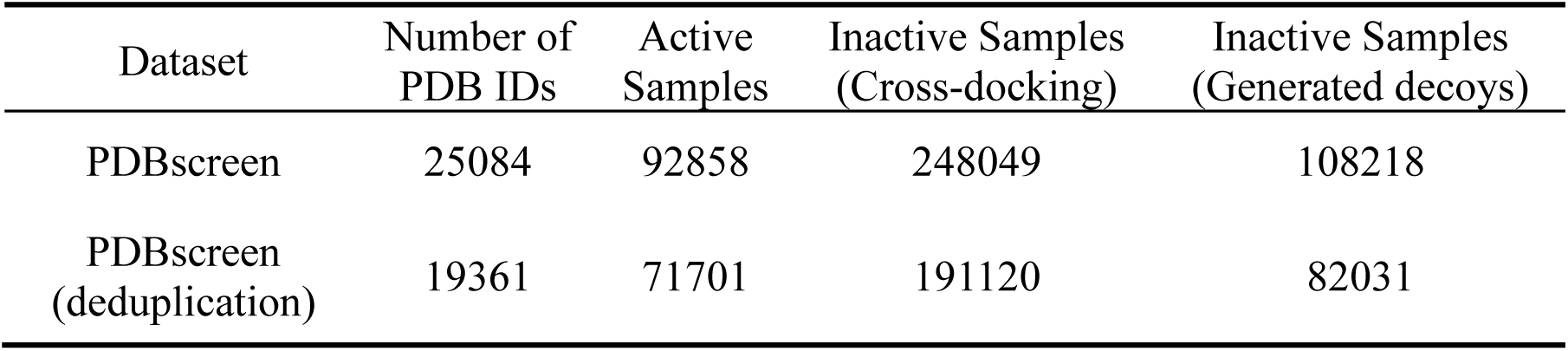
Statistics of PDBscreen.

| Dataset | Number of PDB IDs | Active Samples | Inactive Samples (Cross-docking) | Inactive Samples (Generated decoys) |
| --- | --- | --- | --- | --- |
| PDBscreen | 25084 | 92858 | 248049 | 108218 |
| PDBscreen (deduplication) | 19361 | 71701 | 191120 | 82031 |

### The typical training and testing process has the risk of data leakage

PDBbind, CASF-2016, DUD-E, and DEKOIS2.0 are commonly used datasets for training and testing scoring methods. However, there is “hard overlap” or “soft overlap” data in these databases^17^, which may lead to data leakage and overestimate the method’s generalization performance. For example, there are many scoring methods that have been trained on the PDBbind dataset, and “externally” tested on CASF-2016, DUD-E and/or DEKOIS2.0^13–16, 40^. Proteins that appear in both the training set and the test set were usually remained or simply deduplicated based on their PDB IDs. However, this data preparation scheme may result in the presence of identical proteins in both the training and testing sets, i.e. “soft overlap”^17^, and their bound ligands may share high similarity or similar scaffolds, leading to significant data leakage issues. Here, in **Supplementary Fig. 1**, we firstly analyzed the overlapping data issue between CASF-2016 and PDBbind2020. We found that there are a total of 67 proteins (with unique UniPort IDs) in CASF-2016, all of which have been included in the PDBbind2020 dataset, corresponding to a total of 4471 different complexes (with unique PDB IDs). Using a ligand similarity threshold of 0.5, we found that over 70% (203/285) of the ligands in CASF-2016 have structural analogs that bind to the same protein in PDBbind2020 (after deduplicating with CASF-2016 by PDB IDs). This potential data leakage issue can lead to an overestimation of performance metrics when using the CASF-2016 to evaluate the models trained on the PDBbind2020. In **Table 2**, we summarized the number of overlapping data among commonly used datasets. It can be observed that there are similar situations for DEKOIS2.0 and DUD-E (**Supplementary Fig. 1**). Therefore, we believe that a more rigorous deduplication method must be used to better evaluate the performance of a scoring method, especially its generalization ability to proteins and ligands not seen in the training set.

**Table 2.** Statistics of PDBbind2020, CASF-2016, DUD-E, and DEKOIS2.0.

| Dataset | Number of PDB IDs | Number of Uniport IDs | Number of duplicated Uniport IDs in PDBbind2020 | Number of PDB IDs with duplicated Uniport IDs in PDBbind2020 |
| --- | --- | --- | --- | --- |
| PDBbind2020 | 19443 | 3973 | - | - |
| CASF-2016 | 285 | 67 | 67 | 4471 |
| DEKOIS2.0 | 81 | 77 | 68 | 2433 |
| DUD-E | 102 | 100 | 89 | 4097 |

In this study, we evaluated the generalization ability for unseen targets using DUD-E and DEKOIS2.0 as external test sets. Firstly, we removed data from the training data with the same UniPort ID as the proteins in these two datasets (data statistics after deduplication are shown in **Table 1**). We then divided the training/validation data by Uniport IDs. **Table 1** and **Table 2** show that although we collected more data from the PDB database, EquiScore used fewer complexes for training than PDBbind2020 due to the deduplication.

### Architecture of EquiScore

EquiScore is a binary classification model that assesses the binding potential between a protein and a ligand by inputting the heterogeneous graph constructed by the protein pocket region and the ligand. **Fig. 2** illustrates the architecture of EquiScore. The first step involves constructing a heterogeneous graph with protein pocket and ligand. The second step initializes the representation of the graph’s nodes and edges through their corresponding embedding layer. The third step involves sending the initialized graph to the EquiScore layer to learn its representation. Finally, in task layer, the atomic representation on the ligand is read out, and the output score of the multi-layer perceptron is used for downstream tasks.

**Fig. 2.**
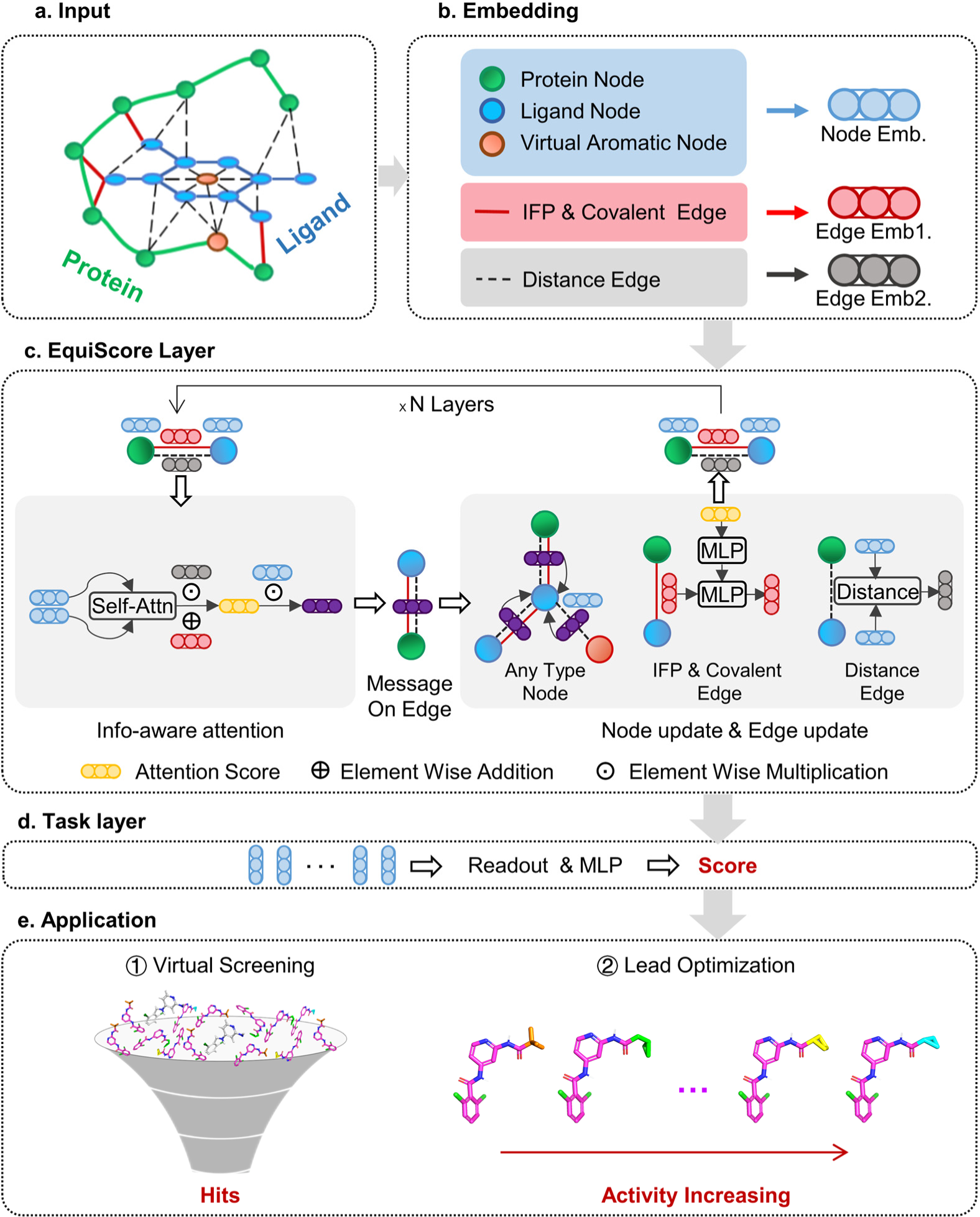
**The overall architecture of EquiScore. a**: Constructing a heterogeneous graph as input. **b**: Embedding layers are used to initialize features into latent space. **c**: EquiScore layers are used for feature extraction and fusion. **d**: ligand’s features are sent to a task layer to predict protein-ligand interaction. **e**: Application scenarios.

In the first step, we designed a heterogeneous graph construction scheme. Aside from abstracting the existing atoms into nodes, we also added a virtual node for each aromatic ring based on expert prior knowledge to better represent the aromatic system. To construct edges, we established geometric distance-based edges (E_geometric_) between nodes and structure-based edges (E_structural_) through chemical bonds. We also added a class of edges in E_structural_ based on protein–ligand empirical interaction components (IFP) calculated by ProLIF^41^ to include prior physical knowledge about intermolecular interaction. In the second step, we used embedding layers to obtain a latent representation for each type of edge and node on the heterogeneous graph.

The EquiScore layer consists of three sub-modules: the info-aware attention module, the node update module, and the edge update module. First, the info-aware attention module uses the distance gating mechanism to model the distance-dependent message passing between atomic pairs. It does this by leveraging the distance information on the E_geometric_ to gate the attention coefficient between atomic pairs. Additionally, the module takes the information on the E_structural_ as the bias item of attention. This allows it to introduce the knowledge of the chemical structure into the model. Second, after obtaining the attention coefficient with geometric and structure information, the info-aware attention module uses it as the coefficient of both vector and scaler features of the neighbor node to update the features of the center node. This ensures the equivariance of the network^42^ in the node update module. Third, when learning the information interaction on different edges^43^, the edge update module uses the attention information on the E_geometric_ to update the features of other types of edges. This allows the information in different types of edges to be better integrated and fused with node information for feature fusion. Finally, after representation learning in EquiScore layers, the ligand’s features are sent to a task layer to predict protein-ligand interaction.

### EquiScore shows improved VS capability on unseen proteins

As analyzed above, the VS capability on proteins not seen in the training set is the most important indicator for evaluating the generalization performance of a scoring method in real-world applications. For comparison, we selected different scoring methods as baselines, including 15 from an earlier evaluation^6^, and added six recently reported models: Kdeep^13^, 3D-GNN^13^, PIGNet^13^, TANKBind^40^, RTMScore^14^, and DeepDock^16^. For the models that have been evaluated previously, we directly referred to the performance metrics reported in the original literature^6^. For the methods that have not been evaluated, we listed the results calculated using their officially reported code and weights. As discussed above, all the previously evaluated methods are established based on the PDBbind dataset, which has a high level of “soft overlap” with the external test sets. To check whether such data leakage would lead to overestimated performance, we removed those samples with proteins that had already appeared in training set for these methods (The target information after deduplication is shown in **Supplementary Table 1** and **Supplementary Table 2**) and added an asterisk annotation to model names to distinguish the new evaluation results (**Fig. 3** and **Fig. 4**).

**Fig. 3.**
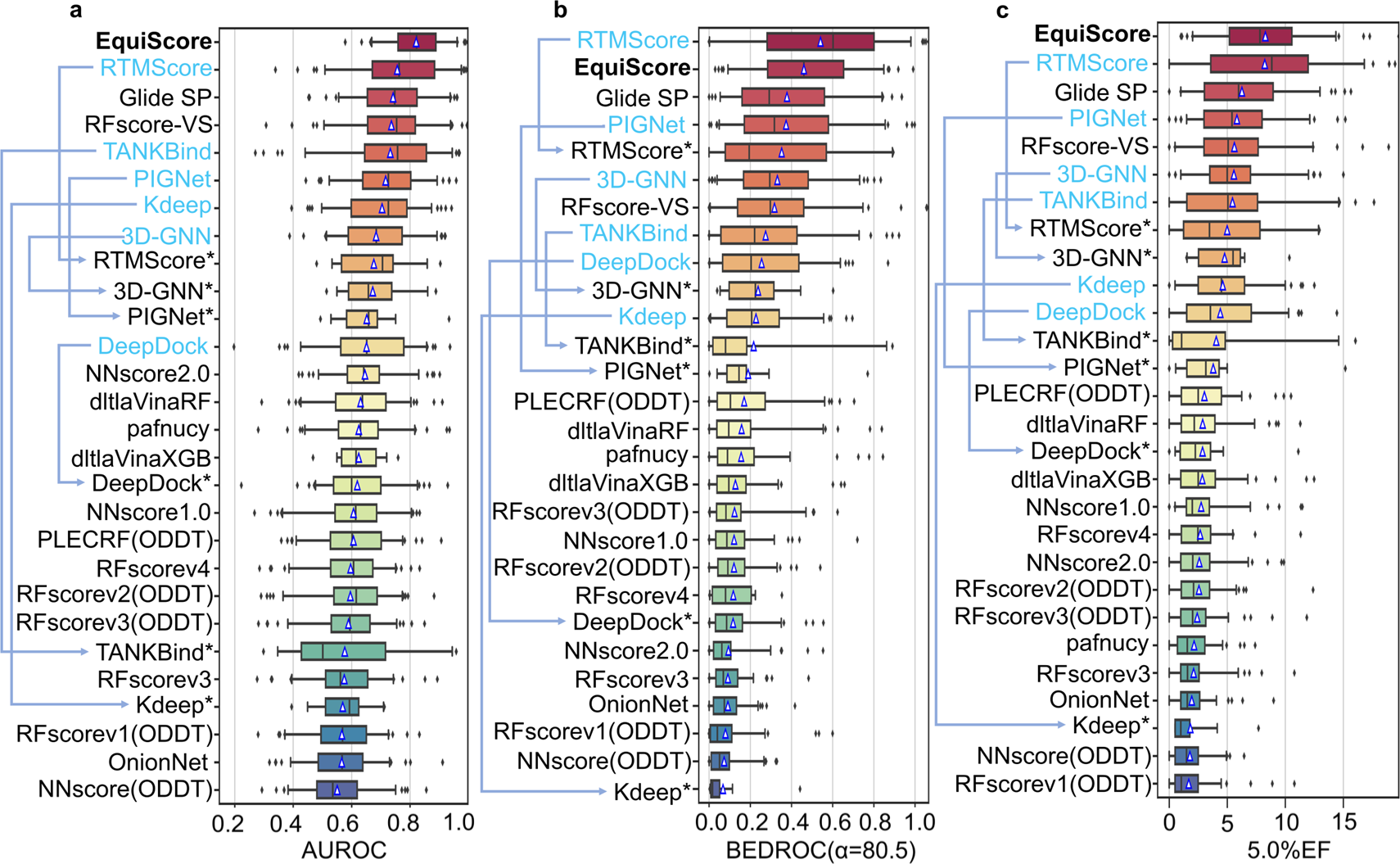
**Evaluation of 22 scoring methods on DEKOIS2.0 in terms of a**: AUROC, **b**: BEDROC (α = 80.5) and **c**: 5.0% EF. The blue triangles in the boxplots represent the means for each bin. All methods are sorted by their mean value. The performance before and after deduplication are marked with blue highlights and asterisks, respectively. Arrowed lines denote the changes in performance ranking.

**Fig. 4.**
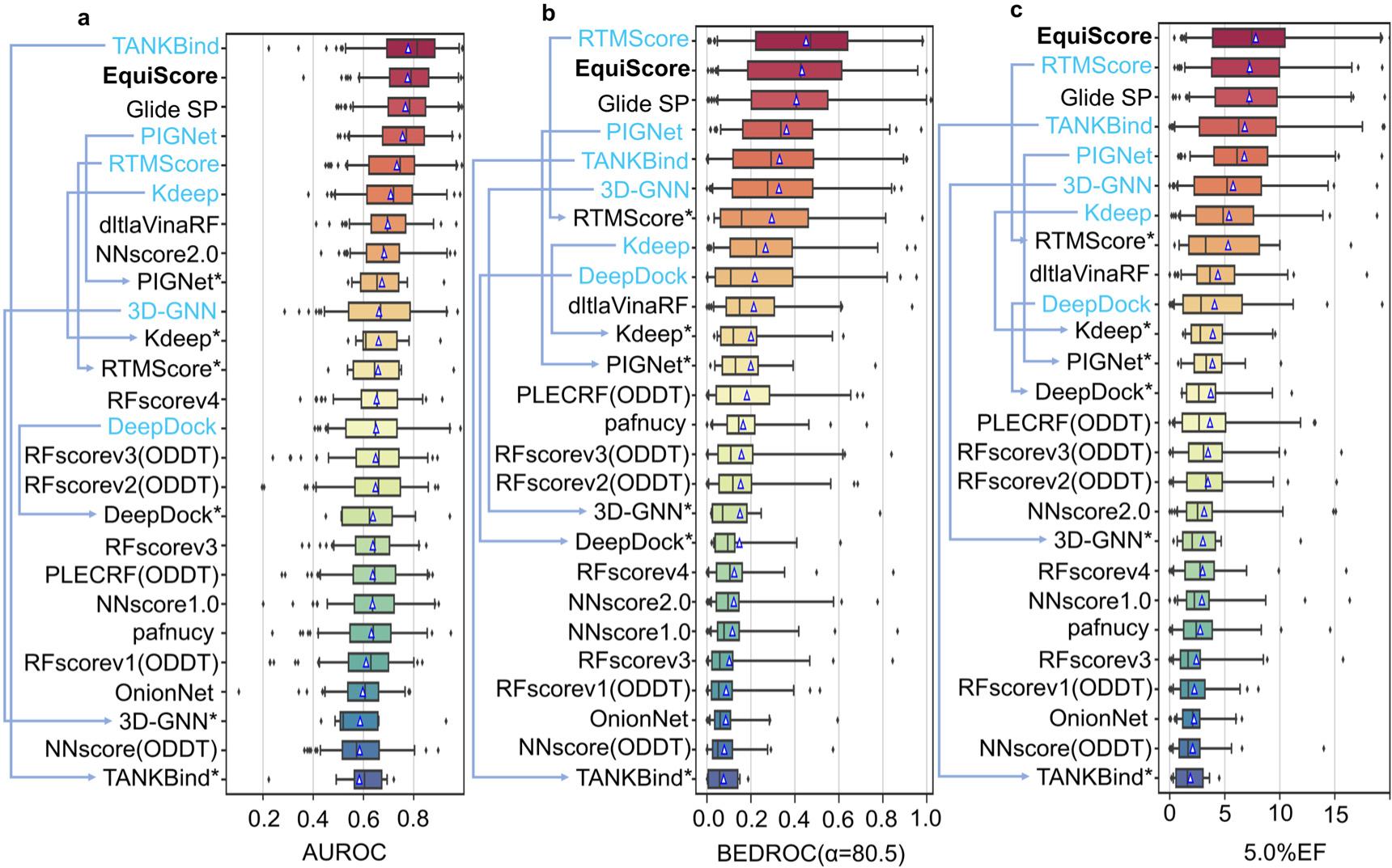
**Evaluation of 22 scoring methods on DUD-E in terms of a**: AUROC, **b**: BEDROC (α = 80.5) and **c**: 5.0% EF. The blue triangles in the boxplots represent the means for each bin. All methods are sorted by their mean value. The performance before and after deduplication are marked with blue highlights and asterisks, respectively. Arrowed lines denote the changes in performance ranking.

We first verified the effectiveness of EquiScore as a rescoring method with the putative binding pose generated via Glide SP. The overall performance was evaluated in terms of area under the receiver operating characteristic curve^44^ (AUROC), Boltzmann-Enhanced Discrimination of ROC^45^ (BEDROC), and enrichment factor^46^ (EF), as shown in **Fig. 3**. EF is defined as the percentage of true binders observed among all of the true binders for a given percentile of the top-ranked candidates (0.5%, 1.0%, or 5.0%) of a chemical library^6^. The BEDROC score considers all compounds rather than a proportion of the chemical library and can be modulated by a parameter α to adjust the weight given to the top-ranked compounds. Here, the α-value was set to 80.5, meaning that the top 2% of ranked molecules accounted for 80% of the BEDROC score^6^.

In **Fig. 3**, we presented the results of our analysis on DEKOIS2.0, which is composed of 81 targets. Each protein has 40 positive compounds extracted from BindingDB^47^ and 1200 decoys generated from ZINC^48^. EquiScore achieved the highest AUROC score of 0.821, which is significantly higher than the second-place RTMscore’s score of 0.756 (**Fig. 3**a). To further compare the early recognition ability of rescoring methods, we calculated and compared the BEDROC metric of all methods shown in **Fig. 3**b. EquiScore outperformed all the baselines except RTMscore and achieved a BEDROC score of 0.460. Remarkably, when considering only the targets not seen during training, the RTMScore performance dropped significantly, from 0.541 to 0.352 (RTMscore*), much lower than EquiScore’s 0.460. In **Fig. 3**, We also observed the same phenomenon in the performance of other methods trained based on PDBbind2020. Regarding EFs, a similar performance drop can be observed when considering both the top 0.5% and 1.0% of ranked compounds (**Supplementary Fig. 2**). When considering the top 5.0% of ranked compounds, EquiScore achieved the highest performance again and had an even greater advantage over other methods when only considering the results on the unseen targets during training. The above results demonstrated that EquiScore’s overall ranking ability significantly exceeds that of existing methods under more rigorous tests. Furthermore, EquiScore’s VS enrichment ability on unseen targets exceeded both traditional scoring methods and deep learning methods^6^.

Subsequently, we extended our evaluation to the DUD-E dataset, which is composed of 102 targets from 8 diverse protein families containing millions of compounds. This allowed us to further verify the screening power of EquiScore in a larger-scale screening scenario. On the DUD-E dataset, TANKBind achieved the best performance on AUROC with a score of 0.778, which has a slight advantage over EquiScore’s 0.776. However, similar to the previous analysis results, we found that the TANKBind’s performance has significantly decreased to 0.583 (TANKBind*) for unseen targets during the training process, even dropping from first to last place. On the other two metrics, BEDROC and 5.0%EF, TANKBind* also showed the worst performance, indicating risk of overfitting during training. Overall, the results on DUD-E were consistent with those of DEKOIS2.0. Other methods trained based on PDBbind2020 also show significant drops in performance on AUROC, BEDROC, and EFs when only considering the unseen targets. In contrast, EquiScore demonstrated advantages over other methods on unseen targets with BEDROC, 1.0%EF, and 5.0%EF scores of 0.432, 17.675, and 7.819, respectively (see **Fig. 4** and **Supplementary Fig. 3**).

### EquiScore shows activity ranking capability on homologous compounds

In the high-throughput VS scenario, a good scoring method must distinguish active molecules from a large batch of inactive molecules by ranking active molecules ahead of inactive molecules through scoring. In contrast, lead compound optimization involves active molecules with similar structures or common scaffolds. In this case, a good method must distinguish subtle differences in activity among these structural analogs. Methods with strong VS capabilities may not have decent analog ranking power, and vice versa. Generally, methods with strong analog ranking power require significantly higher computational cost, such as free energy perturbation^35^ (FEP). Currently, very few methods that can simultaneously demonstrate good VS and analog ranking capabilities while lacking rigorous external validation. To further verify the potential of EquiScore in lead compound optimization scenarios, we collected a dataset containing eight groups of homologues and their activity data from the literature^35^ to test the ranking capability of EquiScore. We named this dataset LeadOpt. For each group of analogs, we computed the scores based on the provided protein-ligand complex structures and compared EquiScore with different methods in terms of Spearman correlation coefficients between the corresponding scores and the activity values. As previously reported^35^, we averaged the total coefficient values weighted by the number of ligands in each group.

To eliminate potential data leakage risks, EquiScore was retrained with the PDBscreen dataset after deduplication based on the Uniport IDs of proteins in LeadOpt. Data statistics after deduplication with LeadOpt are summarized in **Supplementary Table 3**. For the methods that had been previously evaluated, we referred directly to the performance metrics reported in the original literature^35^. For the methods that have not been evaluated, we listed the results calculated using their officially reported weights^13, 14, 16, 40^.

FEP+ is a commercial FEP calculation tool implemented in Schrödinger, which has demonstrated extremely high calculation accuracy in previous report^35^. As shown in Table 4, EquiScore (0.57) ranked second only to FEP+ (0.73) on LeadOpt. This result indicates that EquiScore has ability to distinguish small differences between similar compounds, which is reflected in its ranking performance. While it is still distant from FEP+ in terms of performance, the results are still meaningful as our method is orders of magnitude faster than FEP+.

### EquiScore demonstrates robust rescoring capabilities

As previously mentioned, the EquiScore model was trained based on putative binding poses generated by Glide SP docking. However, deep learning methods are prone to overfitting the training data. Therefore, we investigated EquiScore’s generalizability to poses generated by other docking methods. We collected putative binding poses generated by different docking software (AutoDock Vina, GOLD CHEMPLP, Surflex-Dock, LeDock, Glide SP) on the DEKOIS2.0 dataset and rescored them using EquiScore. Our goal was to determine whether EquiScore, combined with different docking methods, can maintain VS capability. The performance metrics are consistent with those used in previous sections.

**Fig. 5** illustrates the VS performance of each docking method and the comparison after EquiScore rescoring. It is satisfying to notice that EquiScore significantly enhances the VS performance of all docking methods. For the four docking methods with relatively poor performance, three of them can surpass or be comparable to the industry-leading commercial docking method, Glide SP, after EquiScore rescoring. The performance improvement after rescoring on 1% EF is two to three times compared to the original docking methods. EquiScore rescoring can also increase Glide SP’s performance, achieving the highest 1% EF of 16.83 among all combinations.

**Fig. 5.**
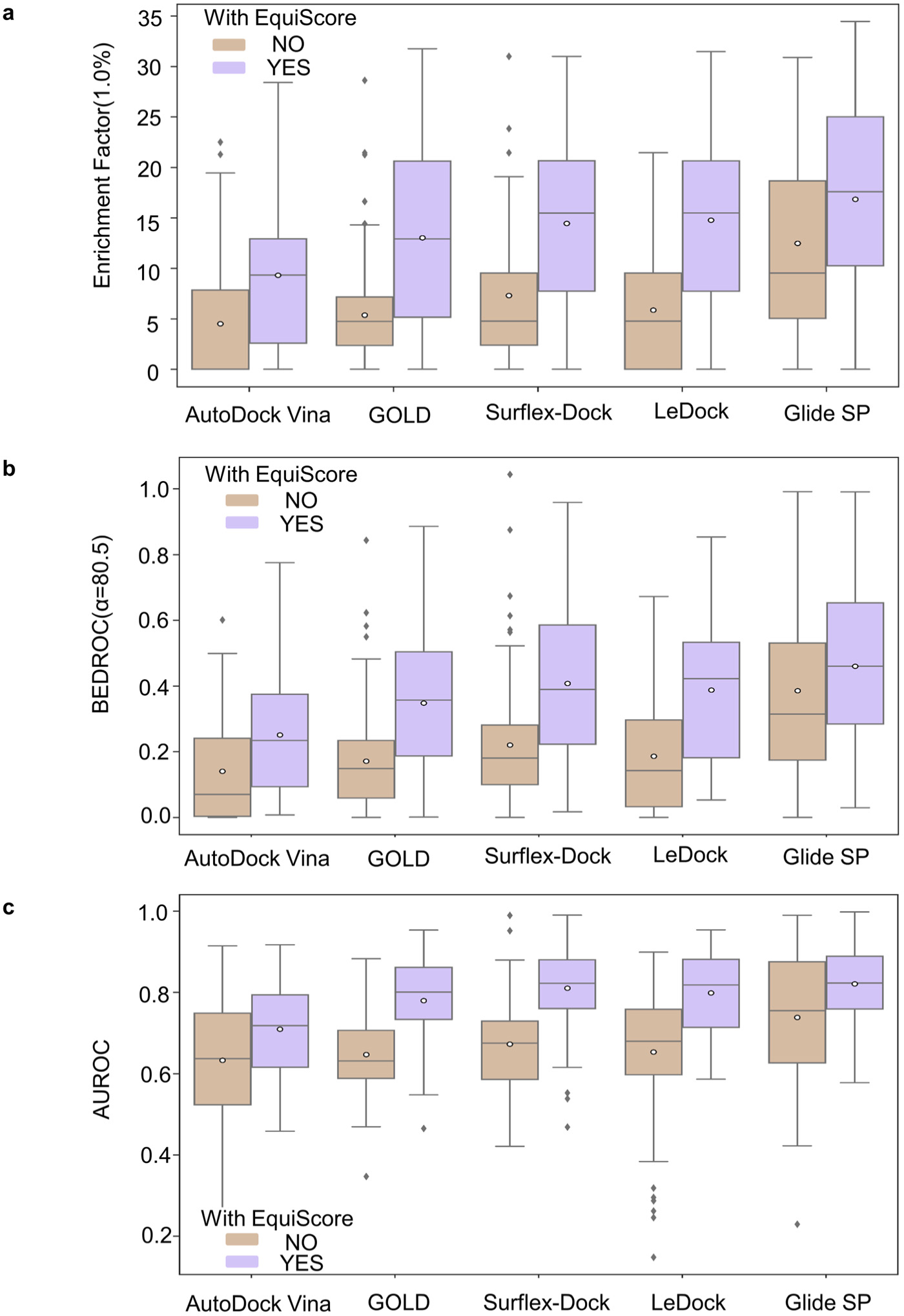
**Performance comparison of EquiScore for rescoring the docking poses generated by different docking methods, in terms of a:** EF (top 1.0%), **b:** BEDROC (α=80.5) and **c:** AUROC.

Unlike EF at 1%, BEDROC and AUROC take all compounds into consideration, rather than just a proportion of the chemical library. Using these two metrics, we may find that EquiScore rescoring improves the screening ability of the original docking methods, and four of them outperform Glide SP after rescoring (see **Fig. 5**b and **Fig. 5**c). Overall, although EquiScore was trained based on putative binding poses generated by Glide SP, it is not sensitive to changes in pose generation during the inference process. This robust rescoring ability may extend the versatility and adaptability of EquiScore, allowing it to seamlessly integrate with various molecular docking methods.

### Ablation study of main strategies

In order to verify the contributions of different modules in Equiscore, we conducted in-depth ablation experiments that ranged from model components to data augmentation strategies.

In the VS scenario, the performance is measured by 1.0% EF on DEKOIS2.0 (**Fig. 6**a). We found that all modules made significant contributions, and removing any of them would lead to performance degradation. Among them, the modules related to data augmentation appeared to be more important and yielded greater effects compared to changes in model architecture. VS is a process to identify a small number of positive samples from a large collection of negative samples. Introducing more challenging negative samples can improve the discriminative ability of the model, thereby reducing the false positive rate and improving the screening enrichment rate. This plays a role similar to generative adversarial learning^49^. When constructing negative samples, we generated decoy molecules that have physical, chemical, and 3D shape patterns similar to active molecules, which forces the model to learn higher-level molecular interaction patterns for correct classification. As a result, we found that this decoy generation strategy indeed had the most prominent performance contribution. The second largest contribution to improving the model is augmenting the positive sample data. This involves introducing near-native poses of active compounds with RMSD < 2Å. As discussed in the previous section, this strategy can increase the model’s robustness. During a VS campaign, it is impossible to access the true binding pose of an active molecule. Therefore, using putative binding poses with noises for modeling can make the model more suitable for real-world applications and significantly improve its performance.

**Fig. 6.**
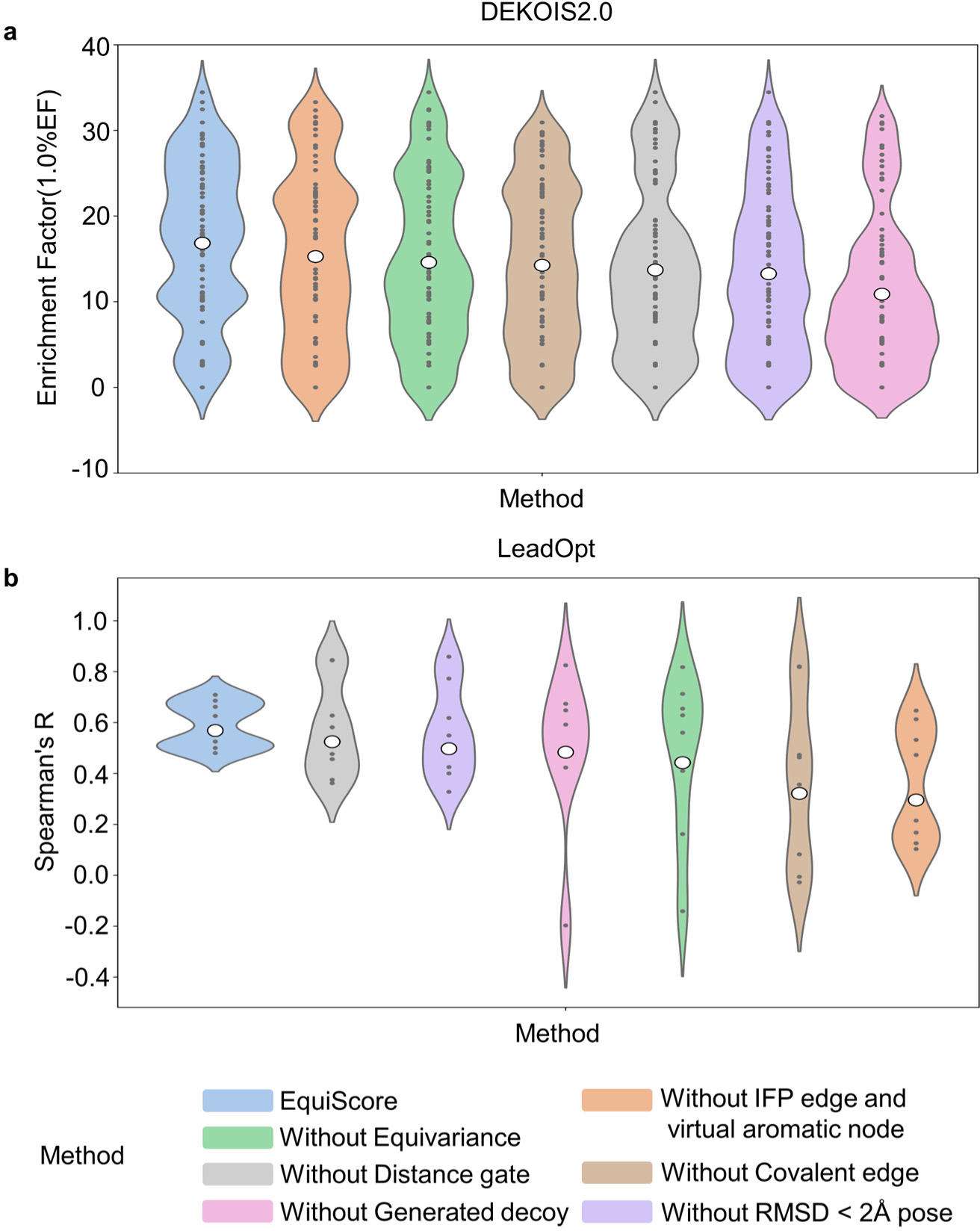
**Ablation results for VS and analogs ranking tasks. a**: VS performance is measured by 1.0% EF on DEKOIS2.0. The white points in the violin plots represent the means for each bin. **b**: Analogs ranking is measured by Spearman’s coefficient on LeadOpt. The white points represent the average of coefficient values weighted by the number of ligands in each group.

In the analogs ranking scenario, the performance is measured by Spearman’s coefficient on LeadOpt (**Fig. 6**b). Similar to previous findings, we discovered that all the modules contributed to the results. Interestingly, in this scenario, the contributions of the modules were completely different from those in VS, and some changes to the model architecture were found more important than data augmentation. We speculate that this may be due to the characteristics of the application. Data augmentation is primarily used to improve the model’s ability to distinguish between positive and negative samples. It may be less suitable for the task of analogs ranking, where most or even all samples are positive. In contrast, the changes in model architecture, particularly the inclusion of physical and chemical knowledge about intermolecular interactions, may enable the model to better capture subtle differences in the interactions between proteins and thus make a more significant contribution to performance.

### EquiScore is interpretable for structure-activity relationship

The EquiScore uses the self-attention module in the Transformer^50^ architecture. Therefore, it is interesting to examine the distribution of attention. Although there is no guarantee that these attention weights are humanly interpretable, in many popular architectures, they do accurately map onto existing concepts^51^. **Fig. 7**a compares the attention weight distribution on IPF edges and covalent edges across 8 attention heads. We can find that all attention heads have different attention weight distributions, suggesting that each head may have learned different components or dependencies of protein-ligand interactions. In our method, IPF edges are constructed based on empirical intermolecular interactions, while covalent edges are built based on covalent bonds. We may notice that the attention weight distributions on these two types of edges are similar in the head 2, but significantly different in the other heads. Multiple heads were originally proposed as a way to address the limited descriptive power of a single head in self-attention^50^. The diversity of attention distributions within and across heads may explain why EquiScore is capable of distinguishing between positive and negative samples, as well as ranking positive samples with close similarity.

**Fig. 7.**
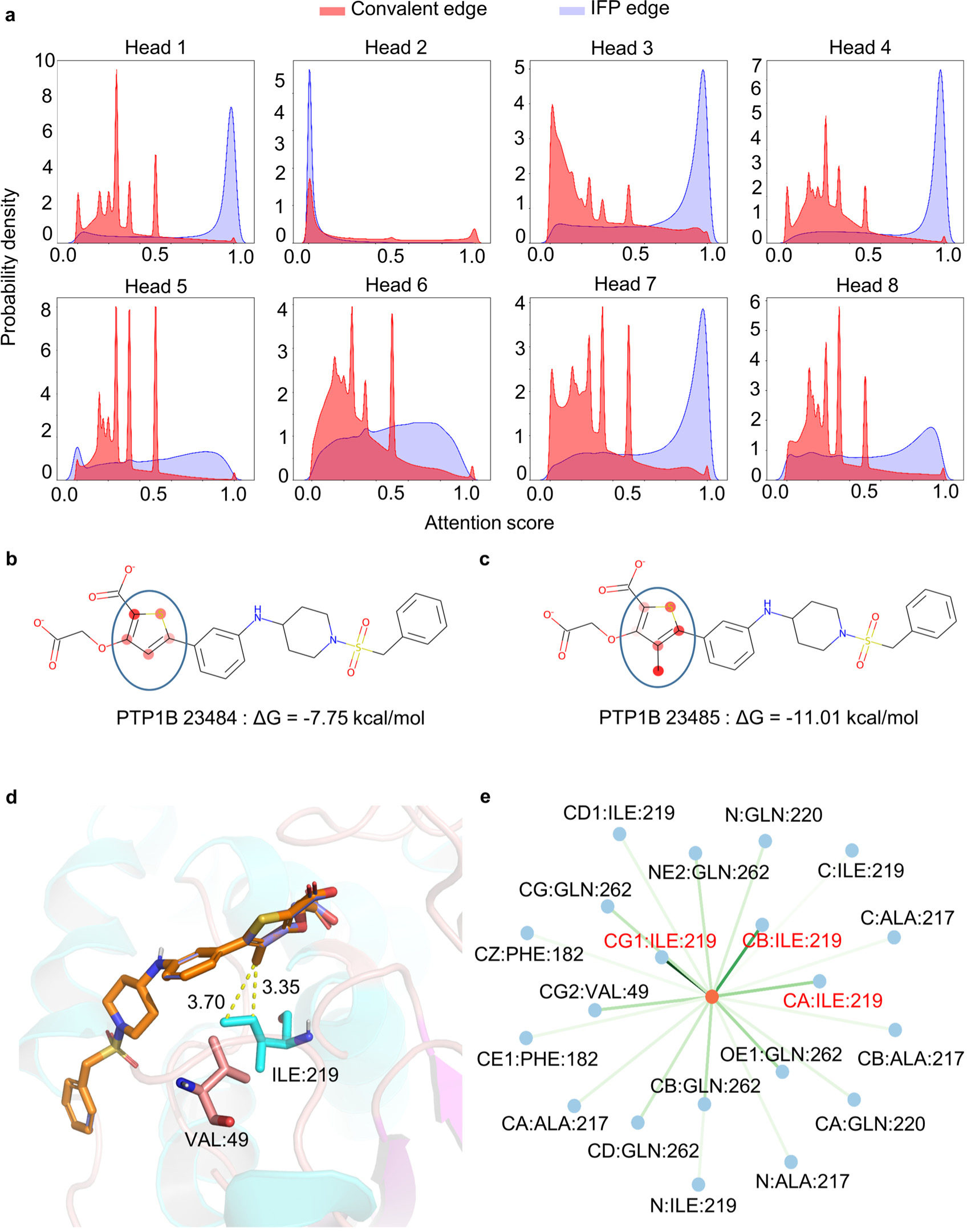
**Interpretation of EquiScore by visualizing attention distribution. a:** Attention score distribution on IFP edges and covalent edges. Attention weights on the ligand **b:** PTP1B 23484 and **c:** PTP1B 23485 (the greater the weight, the darker the color). **d:** The putative binding mode of PTP1B 23485 to human PTP1B (PBD: 2QBS). e: Attention weights of the interaction between the methyl (orange node) of PTP1B 23485 and protein pocket atoms (blue nodes).

In **Fig. 7**b-e, we demonstrate the multi-level interpretability of EquiScore using a lead compound optimization case in LeadOpt. Specifically, in this case, adding a methyl group to the thiophene ring of PTP1B 23484 (**Fig. 7**b) leads to PTP1B 23485 (**Fig. 7**c), which exhibits a “magic methyl effect” ^52^, changing ΔG from −7.7 kcal/mol to −11.01 kcal/mol. By visualizing the attention distribution on the two analogs, we found that the added methyl group was assigned a significantly high weight, indicating its importance. In **Fig. 7**d, we observed that the introduction of the methyl group makes the ligand and the protein pocket more complementary in shape, and brings the molecule and the carbon atoms of the hydrophobic amino acid residues ILE219 and VAL49 in the pocket closer in distance. As shown, the carbon-carbon atom pairs are at a distance of around 3.5Å, which could form favorable hydrophobic interaction^13^. Further visualization of the protein-ligand interaction in **Fig. 7**e reveals that the model assigned a high weight to the introduced methyl with the carbon atoms on ILE219, indicating that the model can well capture ligand atoms and receptors interactions between pairs of atoms through the EquiScore layer. Overall, this interpretability could help us locate key amino acids on protein and optimizable motifs on the ligand, providing a reference for rational drug design.

## CONCLUSION

In this study, we developed a generic protein-ligand interaction scoring method called EquiScore. Firstly, we analyzed common distribution biases and the data leakage issue in developing scoring methods. Based on these analyses, we constructed a new dataset called PDBscreen using multiple data augmentation strategies, such as enlarging the positive sample size with near-native ligand binding poses and the negative sample size with generated highly deceptive decoys. Secondly, leveraging the PDBscreen dataset, we trained a model using an equivariant heterogeneous graph attention architecture that incorporates different physical and prior knowledge about protein-ligand interaction. For example, we defined more types of nodes and edges, including virtual nodes for aromatic rings, spatial distance edges, and empirical molecular interaction edges. Thirdly, we evaluated the performance of the resulting EquiScore model. In the VS scenario, we compared EquiScore with 21 existing scoring methods and found that EquiScore outperformed others on unseen proteins, with the best performance measured by three different metrics: AUROC, BEDROC, EFs scores, on two external datasets DEKOIS2.0 and DUD-E. In the lead compound optimization scenario, we compared EquiScore with eight different types of methods and found that EquiScore showed only lower ranking ability than FEP+. Considering the significantly higher computational expenses required for FEP+ calculations, EquiScore demonstrated the advantage of more balanced speed and accuracy. Additionally, we found that EquiScore demonstrates robust rescoring capabilities when applied to poses generated by different docking methods, and rescoring with EquiScore can enhance the VS performance of all the evaluated methods. Finally, we analyzed the model’s interpretability by studying the self-attention weight distribution of EquiScore and found that the model can capture key inter-molecular interactions, demonstrating the model’s rationality and providing useful clues for rational drug design. Robust prediction of protein-ligand interactions will provide valuable opportunities to learn about the biology of proteins and determine their impact on future drug treatments. We envision that EquiScore may contribute to a greater understanding of human health and disease, and catalyze the discovery of novel medicines.

## METHODS

### Data preparation

#### Data collection and redocking

We followed the process of building a VS database from Adeshina Y O et al.^53^, which involved the following steps: (1) Downloading all protein-ligand complexes in the PDB database (180,207 entries as of July 2021). (2) Retaining crystal structures with resolution better than 2.5 Å. (3) Filtering out complexes with ligand that include nucleotide-like molecules (e.g., ATP), Amina acid-like molecules (e.g., EPW), cofactors (e.g., NAD), crystallographic additive (e.g., polyethylene glycol), or covalent ligand by chemical composition dictionary^54^. The specific pipeline is shown in **Fig.1**.

After redocking the complexes, we kept only the poses with a Root-Mean Square Deviation (RMSD) less than 2Å and the pose with the highest ranking after redocking while keeping a maximum of five poses for each complex. It should be noted that to ensure the data quality of PDBscreen further, we only kept poses with docking scores less than −5kcal/mol in PDBscreen.

#### Docking setup

The optimization of all proteins was performed using the Protein Preparation Wizard of Maestro^55^ module of Schrödinger (version 12.6; Schrödinger, LLC: New York, NY, 2020). This included adding hydrogens, assigning bond orders, filling missing side chains and loops, removing water molecules beyond 5 Å from the ligand, optimizing the H-bond network, and minimizing the system with the OPLS-2005^56^ force field until the root-mean-square deviation of heavy atoms converged to 0.30 Å.

In LigPrep^46^, the Epik^55^ module was used to obtain possible molecular ionization states at a target pH value of 7.0 ± 2.0. The OPLS-2005 force field was then used to generate the ligand conformation with the lowest energy, which served as the starting point for further docking experiments. It is important to note that a single molecule may have multiple tautomers/stereoisomers, which could introduce the analog bias caused by over-represented scaffolds^57^. To mitigate this issue, only the isomers with the best docking scores were saved for further analyses. This means that each molecule with a unique identity had only one docking pose.

The Receptor Grid Generation module generated the receptor grids in Schrödinger with the size of the binding box set to 10 × 10 × 10 Å centered on the co-crystallized ligand. Finally, all the ligand-protein docking was performed with the Glide^46^ module (version 8.9) within Schrödinger using the standard precision (SP) mode.

#### Cross-docking

We use the Uniport^58^ website to map Uniport IDs with PDB IDs in PDBscreen. Uniport is a protein database that contains protein sequences and functional information. Using Uniport ID as a protein identity, the cross-docking process can ensure that the ligands in a PDB ID cannot be docked to related PDB IDs of the same protein. It should be noted that PDBscreen will have the same small molecules under different proteins. In the cross-docking process, we convert all the ligands into the canonical SMILES format, and the same SMILES format is considered to be the same ligand, which further avoids the occurrence of false negatives. Finally, negative samples obtained by cross-docking are ten times larger than positive samples.

#### Generated decoys

To expand the chemical space of ligands in complexes, we use the generative model DeepCoy^38^ to generate 500 decoys for each PDB ID and dock these decoys. Finally, we calculate the 3D degree of overlap of the poses after docking with the ligand in the crystal using the Shape Screening module of Schrödinger. We sort them according to the degree of overlap and only keep the top five poses as final generated decoys. These decoys will work together with the protein as negative samples.

#### Data deduplication

To verify the generalization ability of EquiScore, we removed the “soft overlap” data from the training set by using Uniport IDs to match external test data. This means any sample in the training set that can match the same protein in the external sets DUD-E and DEKOIS2.0 will be excluded. This ensures the proteins of the external sets are unseen during model training.

#### Data pre-processing

When constructing a protein-ligand interaction graph, we only considered residues in the range of 8Å around the ligand and treated atoms as nodes on the graph. For edge construction, we established geometric distance-based edges (E_geometric_) between nodes that were less than 5.5 Å apart and regarded chemical bonds as edges (E_structural_). We added a virtual central node to any aromatic ring that appeared in either the ligand or protein, and established virtual edges between the virtual node and the nodes of the ring. Additionally, we calculated IFP by ProLIF^41^ and established IFP edges between pairs of atoms associated with this interaction. We used the average values of nodes on the relevant ring as the virtual node coordinates and features. The IFP edge feature is a learnable vector, and other defined features are shown in **Table 4**.

#### Model

EquiScore accepts protein-ligand binding pose as input and outputs a score value. For the convenience of description, unless otherwise specified, we use italics to represent variables and bold to represent matrices. Let G = (V, E_geometric_, E_structural_) denote a protein-ligand interaction graph (more description in data preparation). Here, V = {v_1_, v_2_, v_3_, …, v_n_}is a set of nodes, where n is the number of the V, E_geometric_ = {e_1_, e_2_, e_3_, …, e_m_} is a set of edges based on geometric distance, where m is the number of the edges in E_geometric_, E_structural_ = {e_1_, e_2_, e_3_, …, e_k_} is a set of edges based on covalent bonds or IFP information, where k is the number of edges in E_structural_. Every node (v_i_) or edge (e_i_) has a vector *h_i_* or *m_i_* to represent its relative information, respectively.

In this paper, we integrate prior knowledge about physical intermolecular interaction into the heterogeneous graph by introducing new types of nodes and edges. It is also feasible to include a portion of it or introduce other types of nodes or edges during the graph construction phase.

#### Graph Neural Network (GNN)

GNNs aim to learn a representation of nodes or graphs. Typically, modern GNNs follow a learning schema that iteratively updates the representation of a node by aggregating representations of its first or higher-order neighbors^59^, which can be described using the Message Passing Neural Networks (MPNN) framework^60^. This method includes message aggregation, update, and read out three steps. In the message passing phase, the hidden states 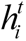 at each node in the graph will update *t* times according to the equation (1) and (2).

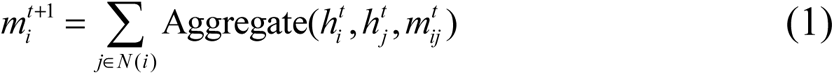

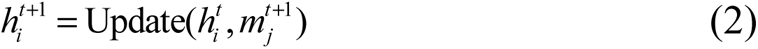

The neighbors of each node and relative edges generate messages and aggregate together by an aggregate function. Then, the node updates to the new hidden state 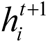 through the aggregated information by the update function. For a graph-level task, a read out function is also required to obtain the representation of the graph. This read out function in the equation (3) must be invariant to permutations of the node states for the MPNN to be invariant to graph isomorphism^60^

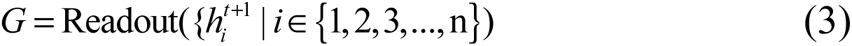

#### Attention mechanism

Our model combines the advantages of graph neural networks and transformer architectures, in which the self-attention module is the main building block.

Let 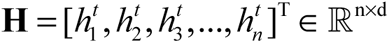 is the input matrix of the self-attention module, d is the dimension of the hidden state 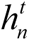, and the calculation process of self-attention with the following equations:

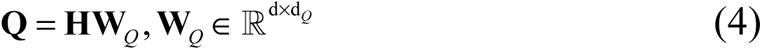

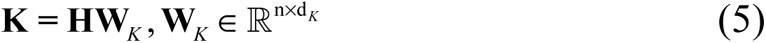

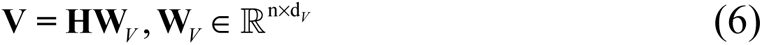

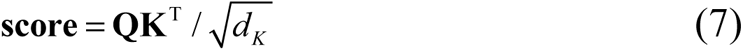

Where **W***_Q_*, **W***_K_*, **W***_V_* are the projection matrices of **H**, the **score**∈ℝ*^n^*^×^*^n^* reflects the similarity of the queries in **Q** and the keys in **K**. For the convenience of expression, we only express the calculation of single-head attention and omit the bias term.

#### EquiScore framework

**(1) Embedding layer:** The features of nodes and edges are mapped to the continuous hidden layer space using a fully connected layer for representation learning.
**(2) EquiScore layer: Fig. 2b.** shows the details for the EquiScore layer. There are three sub-modules in the EquiScore layer: the info-aware attention module, the equivariant message passing module (node update module), and the edge update module.

**(2.1) Info-aware attention module:** This module enables the attention mechanism to aware prior information. After equations (4)-(7), the attention coefficient matrix represents the similarity between each node and other nodes on the graph. However, this method cannot let the model know the 3D structural information on the graph. To use inductive bias from the 3D information to help the model capture distance dependence, the relative distance matrix **E**_geometric_ on geometric-based edges is transformed by the following equation:

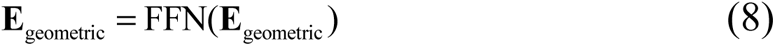

Then, through the equation (9):

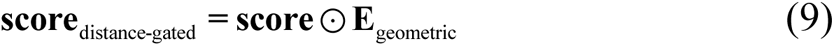

Here, **E**_geometric_ is used as a gating mechanism to control the strength of the information flow of each node to the target node.

To further utilize the information of the geometry-based edges in the graph while preserving the chemical prior information of the complex, an edge bias module is introduced. In this module, the pre-defined feature matrix **E**_structural_ on covalent edges will be a bias term by the following equation:

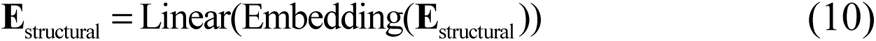

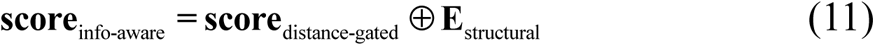

**(2.2) Equivariant message passing module (Node update module):** Following EGNN^42^ ensures the equivariant properties of EquiScore by simultaneously updating the node’s scaler and vector representation using the following equations:

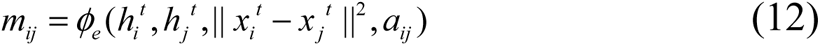

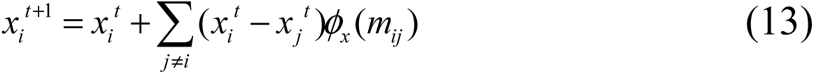

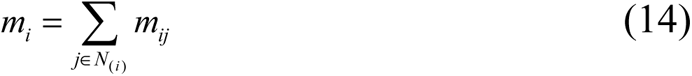

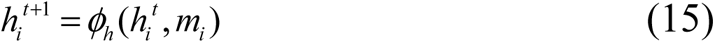

In the equation (12) *a_ij_* is an element in the score matrix **score**_info-aware_, which represents edge features, and *ϕ_e_* is an edge operation function to obtain a massage from input information. In the equation (13), EGNN updates the position of each node as a vector field in a radial direction. In other words, the position of each node is updated by the weighted sum of all relative differences 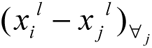, and the weights of this sum are provided as the output of function *ϕ_x_* ^42^. Since our task is an invariant task and positions are static, there is no need to update the atom’s position x. Consequently, we tried both manners and we notice there have some improvement by updating positions, so we kept this step (but only reserve the invariant output to task layer). Finally, equations (14)-(15) follow the same update steps as standard GNNs, like the equation (1) and (2).

**(2.3) Edge-update module:** Many current works^11, 61–63^ also demonstrate that alternate iterative updates of edges and nodes increase the expressiveness and generalization of the model. To further fuse the information carried on different types of edges, we draw inspiration from the work of Dwivedi et al. ^64^ and design an edge update module.

First, we calculate the attention matrix **score**_info-aware_ through equations (8)-(11), and then read out and project the attention matrix to the hidden space with the same dimension as the edge features through a project module consisting of a two-layer neural network and a nonlinear activation function following the equation:

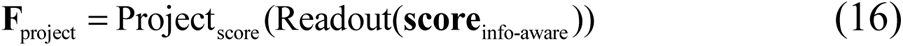

Finally, we add this part of the information through the equation (17) to update the features matrix on structural-based edges.

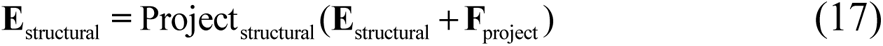

After the above modules, our model acquires rich node and edge representation information, which will be fed into the following parts.

**(3) Task Layer:** After the multi-layer EquiScore layers convolution operation, nodes belonging to the ligand in the ligand-protein interaction graph are read out and fed into a task layer. In this step, we only reserve the invariant output and discard the equivariant output (e.g. updated 3D coordinates) since the goal of this module is to provide invariant features. To ensure permutation invariance, we use the weighted sum operation as the read out pooling function to get the graph-level representation through the following equation:

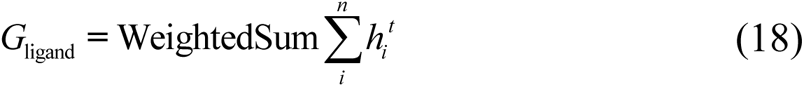

After the graph-level representation *_G_*_ligand_ using the pooling operation, the representation is sent to the multi-layer perceptron layer (MLP_task_) following the equation (19). The multi-layer perceptron layer consists of three linear layers with nonlinear activation function^65^ and finally outputs the predicted probability.

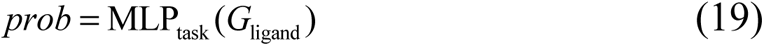

Finally, EquiScore uses cross-entropy as the loss function. Its expression is as the following equation:

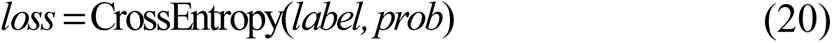

**(4) Model training:** We optimized the model using the Adam^66^ optimizer, with a batch size of 64 and a learning rate of 10e^−4^ without weight decay. The model training proceeded unless the best validation loss did not change in 50 successive epochs.

### External test datasets

#### Lead compound optimization datasets

Schrödinger has reported a lead compound optimization dataset with a broad range of target protein types. The Schrödinger FEP+ workflow was used to calculate the relative binding free energies and correlation coefficients were calculated^35^. To further evaluate the ranking performance of EquiScore and its application potential in lead compound optimization scenarios, we used this dataset, named LeadOpt, as an external test. The statistics of proteins and compounds in LeadOpt are shown in **Table 3**.

**Table 3.** Results on LeadOpt.

| Target | N | Method |  |  |  |  |  |  |  |  |
| --- | --- | --- | --- | --- | --- | --- | --- | --- | --- | --- |
|  |  | DeepDock | PIGNet | Kdeep | 3D-GNN | TANKBind | RTMScore | Glide SP | FEP+ | EquiScore |
| BACE | 36 | -0.13 | -0.20 | 0.42 | 0.41 | -0.19 | -0.05 | 0.11 | 0.74 | 0.53 |
| CDK2 | 16 | 0.11 | -0.41 | 0.84 | 0.80 | 0.73 | -0.42 | -0.36 | 0.41 | 0.66 |
| JNK1 | 21 | 0.62 | 0.70 | 0.45 | -0.51 | 0.41 | 0.66 | 0.27 | 0.90 | 0.69 |
| MCL1 | 42 | 0.49 | 0.13 | 0.45 | 0.21 | 0.70 | 0.45 | 0.50 | 0.78 | 0.48 |
| p38 | 34 | 0.24 | 0.03 | 0.37 | 0.48 | 0.51 | 0.66 | -0.24 | 0.64 | 0.53 |
| PTP1B | 23 | -0.36 | -0.12 | 0.70 | 0.75 | 0.76 | 0.45 | 0.23 | 0.82 | 0.61 |
| Thrombin | 11 | -0.07 | 0.91 | 0.75 | 0.55 | 0.80 | 0.81 | 0.49 | 0.62 | 0.50 |
| Tyk2 | 16 | -0.13 | 0.69 | 0.52 | 0.71 | 0.49 | 0.59 | 0.79 | 0.87 | 0.71 |
| Total | 199 | 0.14 | 0.13 | 0.51 | 0.39 | 0.47 | 0.38 | 0.20 | 0.73 | 0.57 |

**Table 4.** List of Node and Edge Features.

| Node Feature | Description |
| --- | --- |
| Atom from ligand or protein | Identity in pocket or ligand [0, 1] (one-hot) |
| Atom type | C, N, O, S, F, P, Cl, Br, B, H (one-hot) |
| Degree of atom | Number of covalent bonds [0, 1, 2, 3, 4, 5] (one-hot) |
| Hydrogens | Number of connected hydrogens [0, 1, 2, 3, 4] (one-hot) |
| Aromatic | Whether the atom is part of an aromatic ring [0, 1] (one-hot) |
| Formal charge | Electrical charge (integer) |
| Radical electrons | Number of radical electrons (integer) |
| Hybridization | [sp, sp2, sp3, sp3d, sp3d2, other] (one-hot) |
| Chirality | Whether the atom is a chiral center [0, 1] (one-hot) |
| Chirality type | [R, S] (one-hot) |
| Edge Feature | Description |
| Bond type | [single, double, triple, aromatic] (one-hot) |
| Conjugation | Whether the bond is conjugated [0/1] (one-hot) |
| Ring | Whether the bond is in a ring [0/1] (one-hot) |
| Stereo | [StereoNone, StereoAny, StereoZ, StereoE] (one-hot) |

#### Virtual screening datasets

To evaluate the model in different VS scenarios, we chose two benchmark datasets DEKOIS2.0^34^ and DUD-E^20^. DUD-E contains a total of 22886 positive ligands against 102 targets from 8 diverse protein families, including 5 G-protein-coupled receptors (GPCRs), 26 kinases, 11 nuclear receptors, 15 proteases, 2 ion channels, 2 cytochrome P450s, 36 other enzymes, and 5 miscellaneous proteins. These positive compounds were originally retrieved from the ChEMBL09 database^67^. For each positive compound, 50 decoys with similar physicochemical properties but dissimilar 2D topology were generated from ZINC^48^. DEKOIS 2.0 contains 81 structurally diverse targets, each target has 40 positive compounds extracted from BindingDB^47^ and 1200 decoys generated from ZINC. The details of these datasets are available in references^6^.

#### Data availability

The PDBscreen dataset supporting this study’s findings is available in Zenodo with the identifier https://doi.org/10.5281/zenodo.8049380.

The test dataset supporting this study’s findings is available in Zenodo with the identifier https://doi.org/10.5281/zenodo.8047224.

Preprocessed data are provided in this paper.

#### Code availability

The code used to generate the results shown in this study is available under an MIT Licence in the repository https://github.com/CAODH/EquiScore.

## Supporting information

EquiScore supplemental materials

## ACKNOWLEDGMENTS

We gratefully acknowledge financial support from National Natural Science Foundation of China (T2225002 and 82273855), Lingang Laboratory (LG202102-01-02), National Key Research and Development Program of China (2022YFC3400504), SIMM-SHUTCM Traditional Chinese Medicine Innovation Joint Research Program (E2G805H), the open fund of state key laboratory of Pharmaceutical Biotechnology, Nanjing University, China (KF-202301).

## REFERENCE

1. Jumper, J.; Evans, R.; Pritzel, A.; Green, T.; Figurnov, M.; Ronneberger, O.; Tunyasuvunakool, K.; Bates, R.; Zidek, A.; Potapenko, A.; Bridgland, A.; Meyer, C.; Kohl, S. A. A.; Ballard, A. J.; Cowie, A.; Romera-Paredes, B.; Nikolov, S.; Jain, R.; Adler, J.; Back, T.; Petersen, S.; Reiman, D.; Clancy, E.; Zielinski, M.; Steinegger, M.; Pacholska, M.; Berghammer, T.; Bodenstein, S.; Silver, D.; Vinyals, O.; Senior, A. W.; Kavukcuoglu, K.; Kohli, P.; Hassabis, D., Highly accurate protein structure prediction with AlphaFold. Nature 2021, 596 (7873), 583-+.

2. Baek, M.; DiMaio, F.; Anishchenko, I.; Dauparas, J.; Ovchinnikov, S.; Lee, G. R.; Wang, J.; Cong, Q.; Kinch, L. N.; Schaeffer, R. D.; Millan, C.; Park, H.; Adams, C.; Glassman, C. R.; DeGiovanni, A.; Pereira, J. H.; Rodrigues, A. V.; van Dijk, A. A.; Ebrecht, A. C.; Opperman, D. J.; Sagmeister, T.; Buhlheller, C.; Pavkov-Keller, T.; Rathinaswamy, M. K.; Dalwadi, U.; Yip, C. K.; Burke, J. E.; Garcia, K. C.; Grishin, N. V.; Adams, P. D.; Read, R. J.; Baker, D., Accurate prediction of protein structures and interactions using a three-track neural network. Science 2021, 373 (6557), 871-+.

3. Muller, S.; Ackloo, S.; Al Chawaf, A.; Al-Lazikani, B.; Antolin, A.; Baell, J. B.; Beck, H.; Beedie, S.; Betz, U. A. K.; Bezerra, G. A.; Brennan, P. E.; Brown, D.; Brown, P. J.; Bullock, A. N.; Carter, A. J.; Chaikuad, A.; Chaineau, M.; Ciulli, A.; Collins, I.; Dreher, J.; Drewry, D.; Edfeldt, K.; Edwards, A. M.; Egner, U.; Frye, S. V.; Fuchs, S. M.; Hall, M. D.; Hartung, I. V.; Hillisch, A.; Hitchcock, S. H.; Homan, E.; Kannan, N.; Kiefer, J. R.; Knapp, S.; Kostic, M.; Kubicek, S.; Leach, A. R.; Lindemann, S.; Marsden, B. D.; Matsui, H.; Meier, J. L.; Merk, D.; Michel, M.; Morgan, M. R.; Mueller-Fahrnow, A.; Owen, D. R.; Perry, B. G.; Rosenberg, S. H.; Saikatendu, K. S.; Schapira, M.; Scholten, C.; Sharma, S.; Simeonov, A.; Sundstrom, M.; Superti-Furga, G.; Todd, M. H.; Tredup, C.; Vedadi, M.; von Delft, F.; Willson, T. M.; Winter, G. E.; Workman, P.; Arrowsmith, C. H., Target 2035 - update on the quest for a probe for every protein. RSC Med Chem 2022, 13 (1), 13–21.

4. Kaplan, A. L.; Confair, D. N.; Kim, K.; Barros-Álvarez, X.; Rodriguiz, R. M.; Yang, Y.; Kweon, O. S.; Che, T.; McCorvy, J. D.; Kamber, D. N.; Phelan, J. P.; Martins, L. C.; Pogorelov, V. M.; DiBerto, J. F.; Slocum, S. T.; Huang, X.-P.; Kumar, J. M.; Robertson, M. J.; Panova, O.; Seven, A. B.; Wetsel, A. Q.; Wetsel, W. C.; Irwin, J. J.; Skiniotis, G.; Shoichet, B. K.; Roth, B. L.; Ellman, J. A., Bespoke library docking for 5-HT2A receptor agonists with antidepressant activity. Nature 2022, 610 (7932), 582–591.

5. Lyu, J.; Wang, S.; Balius, T. E.; Singh, I.; Levit, A.; Moroz, Y. S.; O’Meara, M. J.; Che, T.; Algaa, E.; Tolmachova, K.; Tolmachev, A. A.; Shoichet, B. K.; Roth, B. L.; Irwin, J. J., Ultra-large library docking for discovering new chemotypes. Nature 2019, 566 (7743), 224-+.

6. Shen, C.; Hu, Y.; Wang, Z.; Zhang, X.; Pang, J.; Wang, G.; Zhong, H.; Xu, L.; Cao, D.; Hou, T., Beware of the generic machine learning-based scoring functions in structure-based virtual screening. Brief Bioinform 2021, 22 (3).

7. Guedes, I. A.; Pereira, F. S. S.; Dardenne, L. E., Empirical Scoring Functions for Structure-Based Virtual Screening: Applications, Critical Aspects, and Challenges. Frontiers in Pharmacology 2018, 9, 1089.

8. Shen, C.; Weng, G.; Zhang, X.; Leung, E. L.; Yao, X.; Pang, J.; Chai, X.; Li, D.; Wang, E.; Cao, D.; Hou, T., Accuracy or novelty: what can we gain from target-specific machine-learning-based scoring functions in virtual screening? Brief Bioinform 2021, 22 (5).

9. Francoeur, P. G.; Masuda, T.; Sunseri, J.; Jia, A.; Iovanisci, R. B.; Snyder, I.; Koes, D. R., Three-Dimensional Convolutional Neural Networks and a Cross-Docked Data Set for Structure-Based Drug Design. J Chem Inf Model 2020, 60 (9), 4200–4215.

10. Ragoza, M.; Hochuli, J.; Idrobo, E.; Sunseri, J.; Koes, D. R., Protein-Ligand Scoring with Convolutional Neural Networks. J Chem Inf Model 2017, 57 (4), 942–957.

11. Li, S.; Zhou, J.; Xu, T.; Huang, L.; Wang, F.; Xiong, H.; Huang, W.; Dou, D.; Xiong, H., Structure-aware Interactive Graph Neural Networks for the Prediction of Protein-Ligand Binding Affinity. In Proceedings of the 27th ACM SIGKDD Conference on Knowledge Discovery & Data Mining, 2021; pp 975–985.

12. Lim, J.; Ryu, S.; Park, K.; Choe, Y. J.; Ham, J.; Kim, W. Y., Predicting Drug-Target Interaction Using a Novel Graph Neural Network with 3D Structure-Embedded Graph Representation. J. Chem. Inf. Model. 2019, 59 (9), 3981–3988.

13. Moon, S.; Zhung, W.; Yang, S.; Lim, J.; Kim, W. Y., PIGNet: a physics-informed deep learning model toward generalized drug-target interaction predictions. Chem Sci 2022, 13 (13), 3661–3673.

14. Shen, C.; Zhang, X.; Deng, Y.; Gao, J.; Wang, D.; Xu, L.; Pan, P.; Hou, T.; Kang, Y., Boosting Protein-Ligand Binding Pose Prediction and Virtual Screening Based on Residue-Atom Distance Likelihood Potential and Graph Transformer. J Med Chem 2022, 65 (15), 10691–10706.

15. Jiang, D.; Hsieh, C. Y.; Wu, Z.; Kang, Y.; Wang, J.; Wang, E.; Liao, B.; Shen, C.; Xu, L.; Wu, J.; Cao, D.; Hou, T., InteractionGraphNet: A Novel and Efficient Deep Graph Representation Learning Framework for Accurate Protein-Ligand Interaction Predictions. J Med Chem 2021, 64 (24), 18209–18232.

16. Méndez-Lucio, O.; Ahmad, M.; del Rio-Chanona, E. A.; Wegner, J. K., A geometric deep learning approach to predict binding conformations of bioactive molecules. *Nat*. Mach. Intell. 2021, 3 (12), 1033–1039.

17. Su, M.; Feng, G.; Liu, Z.; Li, Y.; Wang, R., Tapping on the Black Box: How Is the Scoring Power of a Machine-Learning Scoring Function Dependent on the Training Set? J Chem Inf Model 2020, 60 (3), 1122–1136.

18. Ji, Y.; Zhang, L.; Wu, J.; Wu, B.; Huang, L.-K.; Xu, T.; Rong, Y.; Li, L.; Ren, J.; Xue, D., DrugOOD: Out-of-Distribution (OOD) Dataset Curator and Benchmark for AI-aided Drug Discovery--A Focus on Affinity Prediction Problems with Noise Annotations. arXiv preprint arXiv:2201.09637 2022.

19. Wang, R.; Fang, X.; Lu, Y.; Wang, S., The PDBbind database: collection of binding affinities for protein-ligand complexes with known three-dimensional structures. J Med Chem 2004, 47 (12), 2977–80.

20. Mysinger, M. M.; Carchia, M.; Irwin, J. J.; Shoichet, B. K., Directory of useful decoys, enhanced (DUD-E): better ligands and decoys for better benchmarking. J Med Chem 2012, 55 (14), 6582–94.

21. Scantlebury, J.; Brown, N.; Von Delft, F.; Deane, C. M., Data Set Augmentation Allows Deep Learning-Based Virtual Screening to Better Generalize to Unseen Target Classes and Highlight Important Binding Interactions. J Chem Inf Model 2020, 60 (8), 3722–3730.

22. Morrone, J. A.; Weber, J. K.; Huynh, T.; Luo, H.; Cornell, W. D., Combining Docking Pose Rank and Structure with Deep Learning Improves Protein–Ligand Binding Mode Prediction over a Baseline Docking Approach. J Chem Inf Model 2020, 60 (9), 4170–4179.

23. Chen, L.; Cruz, A.; Ramsey, S.; Dickson, C. J.; Duca, J. S.; Hornak, V.; Koes, D. R.; Kurtzman, T., Hidden bias in the DUD-E dataset leads to misleading performance of deep learning in structure-based virtual screening. PLoS One 2019, 14 (8), e0220113.

24. Pun, G. P. P.; Batra, R.; Ramprasad, R.; Mishin, Y., Physically informed artificial neural networks for atomistic modeling of materials. Nat Commun 2019, 10 (1), 2339.

25. Li, L.; Hoyer, S.; Pederson, R.; Sun, R.; Cubuk, E. D.; Riley, P.; Burke, K., Kohn-Sham Equations as Regularizer: Building Prior Knowledge into Machine-Learned Physics. Phys Rev Lett 2021, 126 (3), 036401.

26. Stärk, H.; Ganea, O.; Pattanaik, L.; Barzilay, R.; Jaakkola, T. In Equibind: Geometric deep learning for drug binding structure prediction, International Conference on Machine Learning, PMLR: 2022; pp 20503-20521.

27. Batzner, S.; Musaelian, A.; Sun, L.; Geiger, M.; Mailoa, J. P.; Kornbluth, M.; Molinari, N.; Smidt, T. E.; Kozinsky, B., E(3)-equivariant graph neural networks for data-efficient and accurate interatomic potentials. Nat Commun 2022, 13 (1), 2453.

28. Zhou, G.; Gao, Z.; Ding, Q.; Zheng, H.; Xu, H.; Wei, Z.; Zhang, L.; Ke, G., Uni-mol: A universal 3d molecular representation learning framework. ChemRxiv 2022.

29. Atz, K.; Grisoni, F.; Schneider, G., Geometric deep learning on molecular representations. Nat. Mach. Intell. 2021, 3 (12), 1023–1032.

30. Batool, M.; Ahmad, B.; Choi, S., A Structure-Based Drug Discovery Paradigm. Int J Mol Sci 2019, 20 (11).

31. Thurlemann, M.; Boselt, L.; Riniker, S., Learning Atomic Multipoles: Prediction of the Electrostatic Potential with Equivariant Graph Neural Networks. J Chem Theory Comput 2022, 18 (3), 1701–1710.

32. Imrie, F.; Bradley, A. R.; Deane, C. M., Generating property-matched decoy molecules using deep learning. Bioinformatics 2021, 37 (15), 2134–2141.

33. Vogel, S. M.; Bauer, M. R.; Boeckler, F. M., DEKOIS: Demanding Evaluation Kits for Objective in Silico Screening — A Versatile Tool for Benchmarking Docking Programs and Scoring Functions. J Chem Inf Model 2011, 51 (10), 2650–2665.

34. Bauer, M. R.; Ibrahim, T. M.; Vogel, S. M.; Boeckler, F. M., Evaluation and optimization of virtual screening workflows with DEKOIS 2.0--a public library of challenging docking benchmark sets. J Chem Inf Model 2013, 53 (6), 1447–62.

35. Wang, L.; Wu, Y.; Deng, Y.; Kim, B.; Pierce, L.; Krilov, G.; Lupyan, D.; Robinson, S.; Dahlgren, M. K.; Greenwood, J.; Romero, D. L.; Masse, C.; Knight, J. L.; Steinbrecher, T.; Beuming, T.; Damm, W.; Harder, E.; Sherman, W.; Brewer, M.; Wester, R.; Murcko, M.; Frye, L.; Farid, R.; Lin, T.; Mobley, D. L.; Jorgensen, W. L.; Berne, B. J.; Friesner, R. A.; Abel, R., Accurate and reliable prediction of relative ligand binding potency in prospective drug discovery by way of a modern free-energy calculation protocol and force field. J Am Chem Soc 2015, 137 (7), 2695–703.

36. Sieg, J.; Flachsenberg, F.; Rarey, M., In Need of Bias Control: Evaluating Chemical Data for Machine Learning in Structure-Based Virtual Screening. J Chem Inf Model 2019, 59 (3), 947–961.

37. Chen, L.; Tan, X.; Wang, D.; Zhong, F.; Liu, X.; Yang, T.; Luo, X.; Chen, K.; Jiang, H.; Zheng, M., TransformerCPI: improving compound-protein interaction prediction by sequence-based deep learning with self-attention mechanism and label reversal experiments. Bioinformatics 2020, 36 (16), 4406–4414.

38. Imrie, F.; Bradley, A. R.; Deane, C. M., Generating Property-Matched Decoy Molecules Using Deep Learning. Bioinformatics 2021.

39. Sastry, G. M.; Dixon, S. L.; Sherman, W., Rapid shape-based ligand alignment and virtual screening method based on atom/feature-pair similarities and volume overlap scoring. J Chem Inf Model 2011, 51 (10), 2455–66.

40. Lu, W.; Wu, Q.; Zhang, J.; Rao, J.; Li, C.; Zheng, S., Tankbind: Trigonometry-aware neural networks for drug-protein binding structure prediction. bioRxiv 2022, 2022-06.

41. Bouysset, C.; Fiorucci, S., ProLIF: a library to encode molecular interactions as fingerprints. J Cheminformatics 2021, 13 (1), 72.

42. Satorras, V. G.; Hoogeboom, E.; Welling, M. In *E (n) equivariant graph neural networks*, International conference on machine learning, PMLR: 2021; pp 9323–9332.

43. Yun, S.; Jeong, M.; Kim, R.; Kang, J.; Kim, H. J., Graph Transformer Networks. Advances in Neural Information Processing Systems 32 (Nips 2019) 2019, 32.

44. Bradley, A. P., The use of the area under the roc curve in the evaluation of machine learning algorithms. Pattern Recognition 1997, 30 (7), 1145–1159.

45. Truchon, J. F.; Bayly, C. I., Evaluating virtual screening methods: good and bad metrics for the “early recognition” problem. J Chem Inf Model 2007, 47 (2), 488–508.

46. Friesner, R. A.; Banks, J. L.; Murphy, R. B.; Halgren, T. A.; Klicic, J. J.; Mainz, D. T.; Repasky, M. P.; Knoll, E. H.; Shelley, M.; Perry, J. K.; Shaw, D. E.; Francis, P.; Shenkin, P. S., Glide: a new approach for rapid, accurate docking and scoring. 1. Method and assessment of docking accuracy. J Med Chem 2004, 47 (7), 1739–49.

47. Liu, T.; Lin, Y.; Wen, X.; Jorissen, R. N.; Gilson, M. K., BindingDB: a web-accessible database of experimentally determined protein-ligand binding affinities. Nucleic Acids Res 2007, 35 (Database issue), D198–201.

48. Irwin, J. J.; Shoichet, B. K., ZINC--a free database of commercially available compounds for virtual screening. J Chem Inf Model 2005, 45 (1), 177–82.

49. Creswell, A.; White, T.; Dumoulin, V.; Arulkumaran, K.; Sengupta, B.; Bharath, A. A., Generative adversarial networks: An overview. IEEE signal processing magazine 2018, 35 (1), 53–65.

50. Vaswani, A.; Shazeer, N.; Parmar, N.; Uszkoreit, J.; Jones, L.; Gomez, A. N.; Kaiser, L.; Polosukhin, I., Attention Is All You Need. Adv Neur In 2017, *30*.

51. Clark, K.; Khandelwal, U.; Levy, O.; Manning, C. D., What does bert look at? an analysis of bert’s attention. arXiv preprint arXiv:1906.04341 2019.

52. Schonherr, H.; Cernak, T., Profound methyl effects in drug discovery and a call for new C-H methylation reactions. Angew Chem Int Ed Engl 2013, 52 (47), 12256–67.

53. Adeshina, Y. O.; Deeds, E. J.; Karanicolas, J., Machine learning classification can reduce false positives in structure-based virtual screening. Proc Natl Acad Sci U S A 2020, 117 (31), 18477–18488.

54. Westbrook, J. D.; Shao, C.; Feng, Z.; Zhuravleva, M.; Velankar, S.; Young, J., The chemical component dictionary: complete descriptions of constituent molecules in experimentally determined 3D macromolecules in the Protein Data Bank. Bioinformatics 2015, 31 (8), 1274–8.

55. Sastry, G. M.; Adzhigirey, M.; Day, T.; Annabhimoju, R.; Sherman, W., Protein and ligand preparation: parameters, protocols, and influence on virtual screening enrichments. J Comput Aided Mol Des 2013, 27 (3), 221–34.

56. Harder, E.; Damm, W.; Maple, J.; Wu, C.; Reboul, M.; Xiang, J. Y.; Wang, L.; Lupyan, D.; Dahlgren, M. K.; Knight, J. L.; Kaus, J. W.; Cerutti, D. S.; Krilov, G.; Jorgensen, W. L.; Abel, R.; Friesner, R. A., OPLS3: A Force Field Providing Broad Coverage of Drug-like Small Molecules and Proteins. J Chem Theory Comput 2016, 12 (1), 281–96.

57. Tuccinardi, T.; Poli, G.; Romboli, V.; Giordano, A.; Martinelli, A., Extensive consensus docking evaluation for ligand pose prediction and virtual screening studies. J Chem Inf Model 2014, 54 (10), 2980–6.

58. UniProt, C., UniProt: the universal protein knowledgebase in 2021. Nucleic Acids Res 2021, 49 (D1), D480–D489.

59. Ying, C.; Cai, T.; Luo, S.; Zheng, S.; Ke, G.; He, D.; Shen, Y.; Liu, T.-Y., Do Transformers Really Perform Bad for Graph Representation? arXiv preprint arXiv:2106.05234 2021.

60. Gilmer, J.; Schoenholz, S. S.; Riley, P. F.; Vinyals, O.; Dahl, G. E., Neural Message Passing for Quantum Chemistry. International Conference on Machine Learning*, Vol* 70 2017, *70*, arXiv:1704.01212.

61. Jiao, Q.; Qiu, Z.; Wang, Y.; Chen, C.; Yang, Z.; Cui, X., Edge-Gated Graph Neural Network for Predicting Protein-Ligand Binding Affinities. In 2021 IEEE International Conference on Bioinformatics and Biomedicine (BIBM), 2021; pp 334–339.

62. Shang, C.; Liu, Q.; Chen, K.-S.; Sun, J.; Lu, J.; Yi, J.; Bi, J., Edge attention-based multi-relational graph convolutional networks. arXiv preprint arXiv:1802.04944 2018.

63. Gong, L.; Cheng, Q. In *Exploiting edge features for graph neural networks*, Proceedings of the IEEE/CVF conference on computer vision and pattern recognition, 2019; pp 9211–9219.

64. Dwivedi, V. P.; Bresson, X., A generalization of transformer networks to graphs. arXiv preprint arXiv:2012.09699 2020.

65. Sharma, S.; Sharma, S.; Athaiya, A., Activation functions in neural networks. Towards Data Sci 2017, 6 (12), 310–316.

66. Xue, Y.; Tong, Y.; Neri, F., An ensemble of differential evolution and Adam for training feed-forward neural networks. Information Sciences 2022, 608, 453–471.

67. Mendez, D.; Gaulton, A.; Bento, A. P.; Chambers, J.; De Veij, M.; Felix, E.; Magarinos, M. P.; Mosquera, J. F.; Mutowo, P.; Nowotka, M.; Gordillo-Maranon, M.; Hunter, F.; Junco, L.; Mugumbate, G.; Rodriguez-Lopez, M.; Atkinson, F.; Bosc, N.; Radoux, C. J.; Segura-Cabrera, A.; Hersey, A.; Leach, A. R., ChEMBL: towards direct deposition of bioassay data. Nucleic Acids Res 2019, 47 (D1), D930–D940.

