## Supplementary material for "EquiScore: A generic protein-ligand interaction scoring method integrating physical prior knowledge with data augmentation modeling": EquiScore supplemental materials

**Corresponding Authors**

### Author Contributions

▽D.H.C., G.C. contributed equally to this study. M.Y.Z. designed the research study. D.H.C developed the method and implemented the code. G.C., collected and processed training data. D.H.C, G.C, J.X.J., D.H.C, and J.Y. benchmarked the methods. All authors contributed to the analysis of the results D.H.C., G. C. and M.Y.Z. wrote the paper. All authors read and approved the manuscript.

### Notes

The authors declare no competing financial interest.

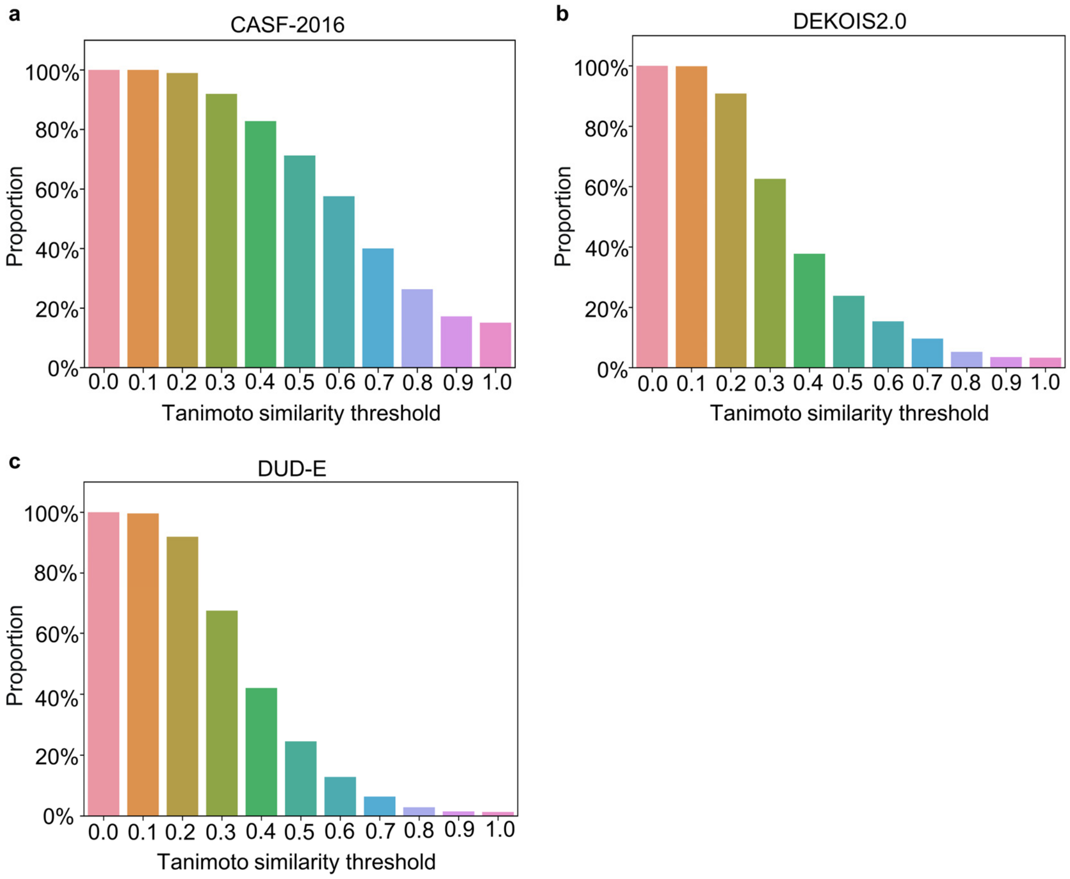


**Supplementary Fig. 1 | Data leakage analysis for CASF-2016, DEKOIS2.0, and DUD-E.** Proportion of active molecules that have “similar” ligands that bind to the same protein in PDBbind2020 (after deduplication with PDB ID) at different similarity thresholds. **a**: CASF-2016, **b**: DEKOIS and **c**: DUD-E.

**Supplementary Table 1 | Information about proteins that did not appear in the PDBbind2020 dataset in DUD-E**

| Target Name | PDB | Classification | UniPort ID |
|---|---|---|---|
| ALDR | 2HV5 | Other Enzymes | P15121 |
| CP2C9 | 1R9O | Cytochrome P450 | P11712 |
| CP3A4 | 3NXU | Cytochrome P450 | P08684 |
| DHI1 | 3FRJ | Other Enzymes | P28845 |
| HXK4 | 3F9M | Other Enzymes | P35557 |
| KITH | 2B8T | Kinase | Q9PPP5 |
| NOS1 | 1QW6 | Other Enzymes | P29476 |
| PGH1 | 2OYU | Other Enzymes | P05979 |
| PPARA | 2P54 | Nuclear Receptor | Q15788 |
| PPARG | 2GTK | Nuclear Receptor | Q15788 |
| PYRD | 1D3G | Other Enzymes | Q02127 |
| SAHH | 1LI4 | Other Enzymes | P23526 |

**Supplementary Table 2 | Information about proteins that did not appear in the PDBbind2020 dataset in DEKOIS2.0**

| Target Name | PDB | Classification | UniPort_ID |
|---|---|---|---|
| 11BETAHSD1 | 3TFQ | Oxido-Reductase | P28845 |
| ACE2 | 1R4L | Protease | Q9BYF1 |
| ALR2 | 1AH3 | Oxido-Reductase | P80276 |
| COX1 | 3KK6 | Oxido-Reductase | P05979 |
| CYP2A6 | 1Z11 | Oxido-Reductase | P11509 |
| ER-BETA | 3OLL | Nuclear Receptor | Q15788 |
| INHA | 1P44 | Oxido-Reductase | P9WGR1 |
| MMP2 | 1HOV | Protease | P08253 |
| PPARA | 2P54 | Nuclear Receptor | Q15788 |
| PPARG | 1FM9 | Nuclear Receptor | Q15788 |
| TK | 1W4R | Kinase | P04183 |

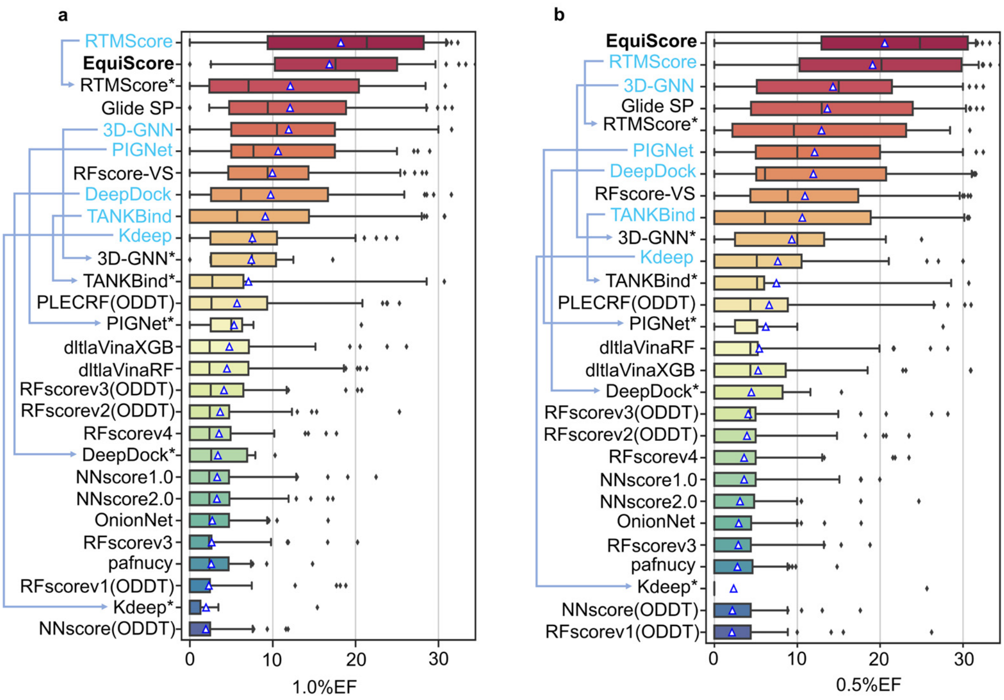


**Supplementary Fig. 2 | Evaluation of 22 scoring methods on DEKOIS2.0 in terms of a**: 1.0% EF and **b**: 0.5% EF. The blue triangles in the boxplots represent the means for each bin. All methods are sorted by their mean value. The performance before and after deduplication are marked with blue highlights and asterisks, respectively. Arrowed lines denote the changes in performance ranking.

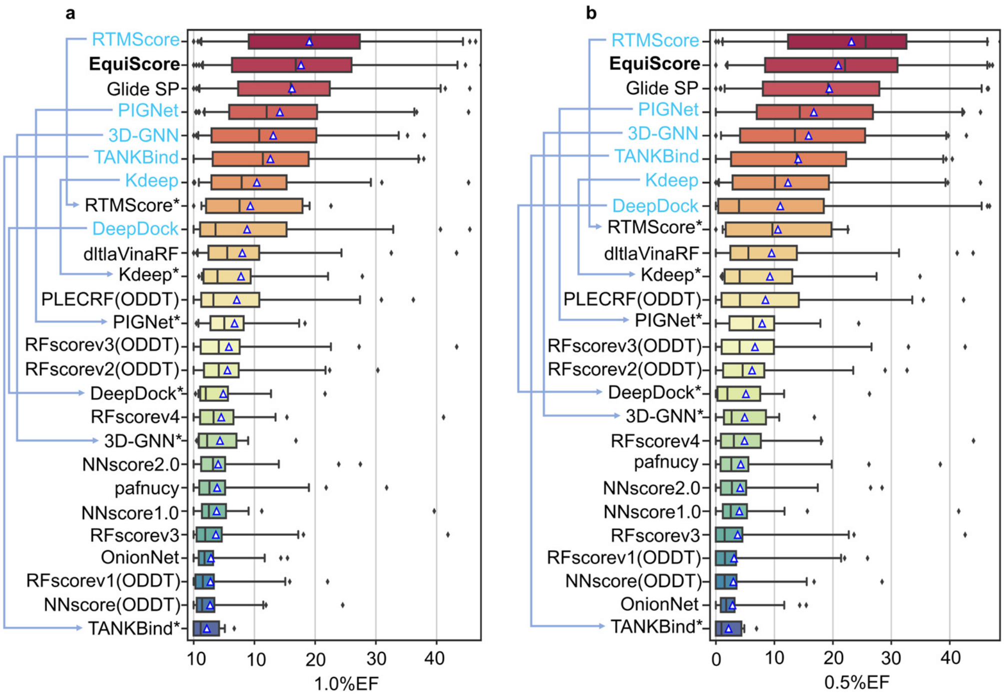


**Supplementary Fig. 3 | Evaluation of 22 scoring methods on DUD-E in terms of a**: 1.0% EF and **b**: 0.5% EF. The blue triangles in the boxplots represent the means for each bin. All methods are sorted by their mean value. The performance before and after deduplication are marked with blue highlights and asterisks, respectively. Arrowed lines denote the changes in performance ranking.

**Supplementary Table 3 | Statistics of data after deduplication with LeadOpt**

| Dataset | Number of PDB ID | Active Samples | Inactive Samples (Cross-docking) | Inactive Samples (Generated Decoys) |
|---|---|---|---|---|
| PDBscreen | 23671 | 87779 | 233955 | 102088 |

**Details of ablation experiments**

To analyze the influence of introducing prior knowledge on the generalization ability of the EquiScore, we conducted ablation experiments as follows.

**(1) Without IFP edge and virtual aromatic node**. In this experiment, we removed the empirical intermolecular interaction edges established by the protein-ligand empirical interaction components (IFP) calculated by ProLIF and virtual aromatic nodes. As seen in **Fig. 6**, the model performance dropped significantly in both two datasets, especially in LeadOpt. We also visualized the attention distribution of covalent and IFP edges in the final EquiScore layer, as shown in **Fig. 7**a. We found that among the eight self-attention heads, the attention weights on IFP edges in seven heads are different from those on covalent edges. This shows that the physical prior knowledge provides additional information for the model, which can effectively help it to improve the generalization ability.

**(2) Without covalent edge.** In this experiment, we removed the information exchange and updated of the two types of edges on the heterogeneous graph in the EquiScore layer. This is equivalent to removing the prior knowledge of the covalent bonding structure in the complex. As shown in **Fig. 6**, the generalization performance of EquiScore dropped significantly in both two datasets, indicating that it is essential to make full use of the original structural prior information and ignore covalent edges will result in the loss of important prior information and lead to poor generalization of the model.

**(3) Without distance gated mechanism.** In this experiment, we removed the distance gating mechanism in the equivariant message passing, which is equivalent to removing the gating effect of the 3D prior information in calculating the attention stage in the EquiScore layer. As in all test results, the model's performance decreased significantly in both datasets, indicating that the 3D distance gating mechanism can effectively enhance the expression ability of the model and better capture the interactions in the complex space. If 3D prior information is only used when constructing the input graph, most information will be lost because some important interactions are distance-dependent. Providing the 3D prior knowledge of distance directly to the model can enable the model to learn this relationship better.

**(4) Without equivariance.** In this experiment, we removed the module that maintains equivariance in the model, and the model degenerated into a standard graph neural network. As seen in **Fig. 6**, the model's performance was greatly affected, indicating that the introduction of the equivariant module enables the model to capture interactions in vector space and scalar space, further increasing the expressive ability of the model.

To investigate the effectiveness of our proposed two data augmentation strategies, we further constructed two ablation experiments.

**(5) Without generated decoy.** As seen in **Fig. 6**, the negative samples formed by the decoys generated by the generative model provide the most significant gain for model generalization in VS scenarios. This result shows that the generated decoys can further help the model learn more complex interaction information instead of learning through some biases of ligands[38].

**(6) Without RMSD < 2Å pose.** For the augmentation strategy of positive samples, the gain of the model in both two datasets is not as high as that of generated decoys, but it still obtains a considerable gain. This result shows that the introduction of diverse poses is also critical to the improvement of model generalization.
